# Dolichol biosynthesis in yeast traces the expanded reaction pathway of higher eukaryotes

**DOI:** 10.64898/2026.04.22.720056

**Authors:** Matthew P. Wilson, Takfarinas Kentache, Charlotte R. Althoff, Isabelle Gerin, Shin-Yi Yu, Marie Morel, Céline Schulz, Sahar Sabry, Jean Jacobs, Julie Graff, Geoffroy de Bettignies, Liliana Surmacz, William C. Sessa, Emile Van Schaftingen, Gert Matthijs, Charles Boone, Kariona A. Grabińska, Guido T. Bommer, François Foulquier

**Affiliations:** Laboratory for Molecular Diagnosis, Center for Human Genetics, KU Leuven, Leuven, Belgium; Metabolic Research Group, de Duve Institute, Université Catholique de Louvain, Brussels, Belgium; Univ. Lille, CNRS, UMR 8576 – UGSF - Unité de Glycobiologie Structurale et Fonctionnelle, Lille, France; Univ. Lille, CNRS, Inserm, CHU Lille, Institut Pasteur de Lille, US 41 - UAR 2014 - PLBS, Lille, France; Biochemical Genetics Department, Human Genetics and Genome Research Institute, National Research Centre (NRC), Cairo, Egypt; Institute of Biochemistry and Biophysics, Polish Academy of Sciences, Pawinskiego 5a, 02-106 Warsaw, Poland; Department of Pharmacology, and Vascular Biology and Therapeutics Program, Yale University School of Medicine, New Haven, Connecticut, 06520, USA; Donnelly Center for Cellular and Biomolecular Research, University of Toronto, Toronto, Canada; Department of Molecular Genetics, University of Toronto, Toronto, Canada; Molecular Ligand Target Research Team, RIKEN Centre for Sustainable Resource Science, Wako, Japan; Cellular and Molecular Physiology Department, Yale School of Medicine, New Haven, CT, USA; Systems Biology Institute, Yale West Campus, West Haven, CT, USA

**Keywords:** N-glycosylation, Yeast, *Saccharomyces cerevisiae*, Congenital Disorders of Glycosylation, Dolichol metabolism, Oxidoreductase, Dolichal, Dolichol, Polyprenol, Polyprenal, Short-chain oxidoreductase, Mannan, Cell wall synthesis

## Abstract

The recent reassignment of *Saccharomyces cerevisiae* Dfg10 as a polyprenal reductase left two unresolved steps in yeast dolichol biosynthesis: the conversion of polyprenol to polyprenal and dolichal to dolichol. In humans, both of these steps are carried out by DHRSX, an enzyme with a unique dual specificity. We found that yeast Env9 catalyzes an NADPH-dependent reduction of dolichal to dolichol. However, in contrast to DHRSX, Env9 does not catalyze the conversion of polyprenol to polyprenal. Instead, our data indicates that this reaction is catalyzed by another yeast short chain oxidoreductase, Tda5. Thus, we provide evidence that the dual role of DHRSX in human dolichol synthesis is fulfilled by two dedicated yeast enzymes, Env9 and Tda5. Accordingly, deletion of *ENV9* and *TDA5* led to the accumulation of polyisoprenoid intermediates, transfer of immature lipid-linked oligosaccharides onto nascent proteins, and defective N-glycosylation. This is similar to what had been observed in DHRSX-deficient mammalian cells and yeast cells with *DFG10* deleted. Furthermore, we discovered that loss of Dfg10 results in a deficiency of cell wall α-mannan, revealing a critical sensitivity of yeast mannan biosynthesis to the quality of nascent N-linked glycans.

## Introduction

Availability of dolichol, a long polyisoprenoid alcohol, is critical for glycosylation, a key post-translational modification in eukaryotic cells. Dolichol phosphates carry mannose and glucose for O/C-mannosylation and GPI anchor synthesis. Dolichol pyrophosphate anchors the lipid-linked oligosaccharide (LLO) to the ER membranes before it is transferred to nascent glycoproteins.^1^

Dolichol exhibits species-specific variation in their isoprene chain length, ranging from Dol-14 to Dol-19 in yeast to Dol-16 to Dol-23 in humans. The first three isoprene units of dolichol are in a *trans*-configuration, the remainder in *cis*-configuration. The terminal unit is saturated, which is crucial for its function as a lipid carrier in glycosylation.^2,3^ Dolichol synthesis is initiated by the heteromeric *cis*-prenyl transferase complex, composed of the NUS1 and DHDDS proteins in humans, which catalyzes the addition of 12–18 isoprene units from isopentenyl diphosphate (IPP) to farnesyl diphosphate (FPP)^4,5^, a precursor molecule that also serves as a substrate for cholesterol and ubiquinol synthesis.^6^ The reaction produces polyprenol pyrophosphate, which is further processed to polyprenol, a compound that still contains a C2-C3 double bond in the last isoprene unit.^7^ Bacteria use a shorter form of polyprenol, undecaprenol, for N-glycosylation.^2^ However, in Eukaryotes this C2-C3 double bond must be reduced to form dolichol for *N*-glycosylation to proceed efficiently.^8,9^ Genetic disorders disrupting dolichol synthesis in humans are categorized as Congenital Disorders of Glycosylation (CDG), with symptoms affecting multiple organ systems.^10–13^

Based on genetic data, it had been assumed that polyprenol is directly converted into dolichol by *H. sapiens* SRD5A3 or the budding yeast *S. cerevisiae* Dfg10.^3,14,15^ However, our recent investigation of a newly identified CDG in patients with pathogenic variants in the *DHRSX* gene allowed us to revise the dolichol biosynthesis pathway. We demonstrated that, in mammals, dolichol formation from polyprenol involves three reaction steps and two polyisoprenoid intermediates.^11,16^ DHRSX first converts polyprenol to polyprenal (**Reaction 1, Fig 1A**), SRD5A3 then reduces this resultant polyprenal to produce dolichal (**Reaction 2, Fig 1A**), which is further reduced to dolichol, again by DHRSX (**Reaction 3, Figure 1A**). In fact, DHRSX was able to use both NAD and NADP as cofactors with almost identical efficiency, likely profiting from a high NAD^+^ to NADH ratio for **Reaction 1** and high NADPH to NADP^+^ ratio for **Reaction 3**. The dual usage of the enzyme DHRSX for an oxidation as well as a reduction reaction in the same pathway was surprising and is, to our knowledge, unique in human metabolism. In multicellular organisms, this dual cofactor usage is compatible with efficient dolichol biosynthesis because the redox states of NAD and NADP are kept relatively stable.^17^

**Figure 1.**
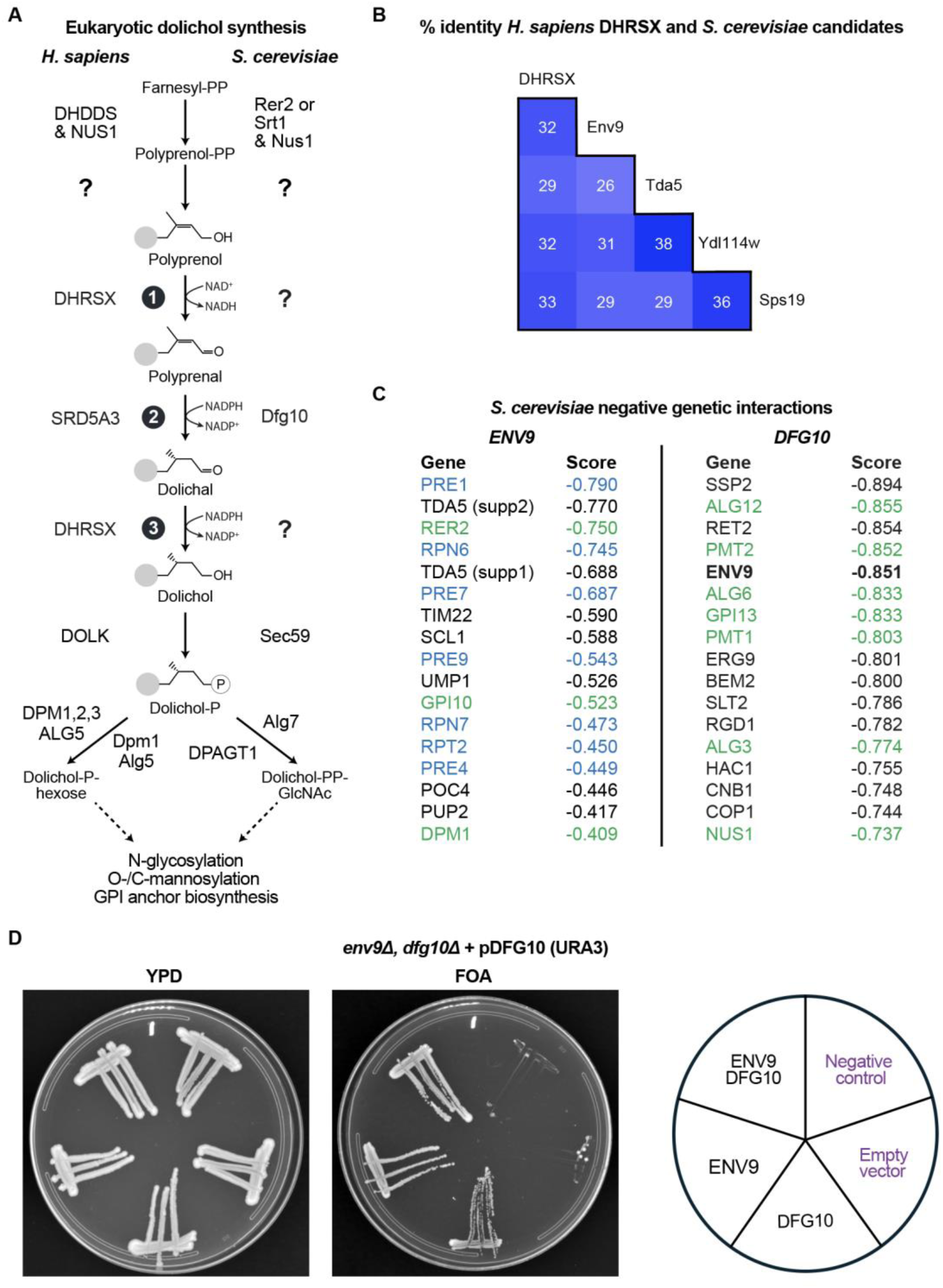
Env9 is a promising *S. cerevisiae* candidate for the functional orthologue *of H. sapiens* DHRSX. (A) The *H. sapiens* (left side) and *S. cerevisiae* (right side) dolichol synthesis pathways. Activities for which a responsible enzyme has not been identified are indicated by ‘?’. Only selected and relevant cofactors are included. Reactions in the pathway of polyprenol to dolichol conversion are numbered 1-3. (B) Protein % identity of *H. sapiens* DHRSX and *S. cerevisiae* Env9, Tda5, Ydl114w and Sps19. See **Table S1** for an extended list of candidate DHRSX orthologs and **Figure S1A** for full alignments of these proteins to DHRSX. (C) A ranked list of the 17 strongest negative genetic interactions for ENV9^24,25^ and DFG10 (this study). Gene mutants that contain deletions or mutations in proteasomal or glycosylation genes are indicated in blue and green, respectively. See **Table S2** and **Table S3** for systematic gene names, mutation alleles and p-values. (D) The *env9Δ dfg10Δ* double deletion strain, expressing *DFG10* from a URA3 plasmid, was transformed with LEU2 single-copy plasmid containing *ENV9* and single-copy plasmid MET17 expressing *DFG10*, or empty vectors. The cells were streaked onto complete (YPD) medium or synthetic complete medium containing 1% 5-fluoroorotic acid (FOA), which selects for cells that have lost the URA3 plasmid. Growth was assessed after ∼2 days (data shown) and monitored for 9 days. Conditions that do not allow growth are indicated in purple. Negative control: yeast strain KG804 transformed with empty vectors.

Our previous work demonstrated that *S. cerevisiae* Dfg10 is a polyprenal reductase and the orthologue of human SRD5A3, revealing that the three-step pathway for polyprenol-to-dolichol conversion is conserved between mammals and yeast.^11^ However, this finding raised a crucial question: given the unstable and highly changeable environment- and nutrition-dependent fluctuations of yeast’s NAD(P) redox state^18,19^, how could a continuous supply of dolichol be ensured? To address this question, we sought to identify and characterize the yeast ortholog of the mammalian enzyme DHRSX. Unexpectedly, we discovered that in *S. cerevisiae*, the dual functions performed by DHRSX in mammals is fulfilled by two separate enzymes, Env9 and Tda5, that might help ensure dolichol synthesis under changing redox conditions. Furthermore, inactivation of these dolichol synthesis genes led to severe, but variable, underglycosylation of nascent glycoproteins, both due to formation of immature N- glycan precursors and their insufficient transfer onto proteins. Finally, we demonstrated that the loss of Dfg10 led to the accumulation of truncated Man_5_GlcNAc_2_ intermediates, compromising formation of mannans, critical structures of the yeast cell wall.

## Results

### Env9, a potential functional orthologue of human DHRSX

To identify enzyme candidates for yeast polyprenol dehydrogenase and dolichal reductase activities, we first performed a BLASTP analysis of *H. sapiens* (h)DHRSX (Uniprot: Q8N5I4) amino acid sequence against the *S. cerevisiae* proteome. This analysis revealed eleven putative oxidoreductase enzymes with E-values <0.01 (**Table S1, Fig S1A).** The *ENV9* (*YOR246C*) product showed the highest degree of similarity, with an E-value of 7×10^-28^ and a protein sequence identity of 27.9% compared to DHRSX. Several additional computational methods listed in the Alliance of Genome Resources^20^ database supported this orthology (https://www.alliancegenome.org/gene/HGNC:18399#orthology).

*S. cerevisiae* Env9 is a poorly characterized short-chain dehydrogenase/reductase (SDR) protein localized to lipid droplets, with a role in maintaining both lipid droplet numbers and size.^21^ It displays weak reductase activity on HMG-CoA and 4-hydroxynonenal *in vitro*^21^, but it is unclear how these activities might be linked to lipid droplet biology. Additional genetic screens have revealed its involvement in vacuolar morphology and lysosomal protein trafficking, suggesting a potential role in intracellular compartmental organization.^22^

Based on the likelihood that Env9 shares a common evolutionary origin and similar function with DHRSX, we used synthetic genetic array (SGA) analysis to search for more evidence implicating it as a candidate enzyme in dolichol synthesis. This technique has mapped negative and positive genetic interactions on a genome-wide scale.^23–25^ Strikingly, *ENV9* showed a strong negative genetic interaction with *RER2* (**Fig 1C, green text**). Rer2, alongside Srt1, is one of the two possible catalytic subunits of the yeast *cis*-prenyltransferase complex. Both these proteins, when in an active complex with Nus1, are responsible for polyprenol formation.^12,26^ A hypomorphic allele is therefore expected to reduce the abundance of polyprenol, the substrate for the last three steps in dolichol synthesis (**Reactions 1-3, Fig 1A**). This observation is consistent with the notion that Env9 acts in the same metabolic pathway as Rer2, which is essential in yeast. *ENV9* also showed strong negative genetic interactions with *GPI10* and *DPM1* (**Fig 1C, green text**), genes involved in dolichol-dependent glycosylation pathways. Furthermore, among the top 17 genes showing the strongest negative genetic interactions, 7 genes (**Fig 1C, blue text**) play a role in proteasomal degradation, a process that helps to eliminate incorrectly folded mis-glycosylated proteins.^27–29^

Many genes included in genome-wide genetic interaction maps have been annotated with the canonical function of their protein product. Therefore, analysis of genetic interactions between genes of known and unknown function can help the formation of hypotheses on the role of uncharacterized genes such as *ENV9*. This process can be visualized by Spatial Analysis of Functional Enrichment (SAFE)^24^, examinable at thecellmap.org.^23^ In this dataset, among the genes showing negative interactions with *ENV9*, genes with annotations for roles in ‘glycosylation’, ‘protein folding’, ‘protein turnover’, and ‘cell wall function’ were most enriched (**Fig S1B**). Taken together, these high-throughput genetic data were consistent with a role of Env9 in dolichol synthesis and a functional contribution to protein glycosylation homeostasis in the ER.

If *ENV9* were implicated in dolichol synthesis, we would also expect a negative interaction with *DFG10*, encoding the yeast polyprenal reductase (**Reaction 2, Fig 1A**). Unfortunately, the high-throughput genetic interaction study did not contain a *bona fide dfg10Δ* strain.^24^ For this reason, we could not immediately interrogate whether there was any genetic interaction between *ENV9* and *DFG10*. To address this, we performed SGA using a newly created *dfg10Δ* strain. Using this strain, *ENV9* (**Fig 1C, bold**) was among the strongest negative genetic interactions identified (**Fig 1C)**. Additionally, 7 genes involved in glycosylation pathways were among the 17 strongest negative interactions with *DFG10* (**Fig 1C, green text**).

In order to confirm this genetic interaction between *ENV9* and *DFG10*, we generated *dfg10Δ env9Δ* double mutant strains carrying a counter-selectable *URA3*-based plasmid expressing both *DFG10*. This strain was complemented with vectors driving expression of both, either of the two genes, or an empty vector (EV) cassette, using *LEU2* and *MET17* selection markers. Upon growth in the presence of 5-fluoroorotic acid (FOA), which selects against *URA3*-based plasmids, we observed that cells were viable in the absence of either *ENV9* or *DFG10*, but the absence of both genes was lethal, confirming that *ENV9* shows an extremely negative genetic interaction with *DFG10* (**Fig 1D**).

In a textbook understanding of a linear metabolic pathway, if one catalytic step is completely blocked, the ablation of another step in the same metabolic pathway should not lead to an increased growth defect. However, even though dolichol biosynthesis in yeast is an essential process, *dfg10Δ* and *env9Δ* deletion mutants are viable. Thus, yeast cells must retain residual dolichol synthesis when either of these major enzymes are absent. Therefore, if either the *dfg10Δ* or *env9Δ* deletion mutant are already relying on imperfect ‘backup’ activities for dolichol formation, then the combined ablation of both main enzymes would likely result in a more severe phenotype, consistent with the observed negative genetic interaction in **Fig 1D**.

### Env9 is a *S. cerevisiae* dolichal reductase

DHRSX-deficient human cells accumulate polyprenol and have reduced dolichol.^11^ We therefore analyzed polyisoprenoids in different yeast strains by LC-MS to understand whether Env9 might indeed be the functional ortholog of DHRSX. In *env9Δ* cells, we observed a strong increase in dolichal (**Fig 2A(v), Fig S2)**, accompanied by a mild reduction in dolichol levels (**Fig 2A(iv), Fig S2)**. Plasmid-based expression of yeast *ENV9* or *DHRSX* partially reversed these changes. We also observed a minor increase in polyprenal (**Fig 2A(ii)**), but it was much less pronounced than in *dfg10Δ* strains. Surprisingly, we did not find any increase in polyprenol (**Fig 2A(i)**). The dolichal accumulation without polyprenol accumulation observed in *env9Δ* yeast cells suggested that Env9 does not perform the two functions associated with DHRSX. Instead, we hypothesized that it may act as a dedicated dolichal reductase (**Reaction 3**, **Fig 1A**).

**Figure 2.**
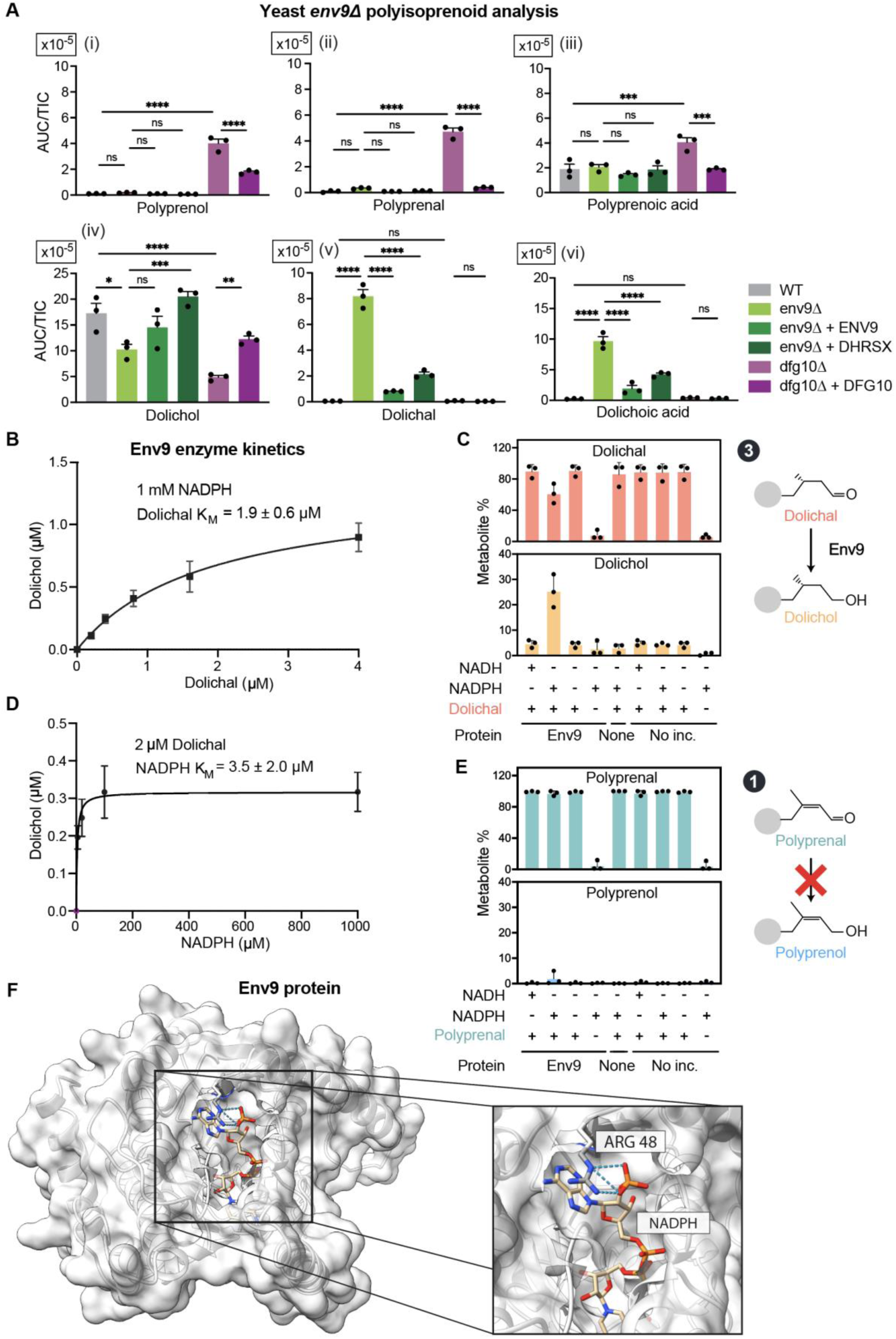
Env9 is a dolichal reductase but not a polyprenol dehydrogenase. (A) Polyisoprenoid species in wild-type (WT), *env9Δ*, and *dfg10Δ* yeast cells and respective complementations. Data represent area under the curve (AUC) normalized to total ion count (TIC) (means ± SEM; *p< 0.05; **p< 0.01; n=3; ***p< 0.001; ****p< 0.0001; n=3). Here and in subsequent figures, the polyisoprenoid species with 16 isoprene repeats is shown. additional chain lengths and chromatograms are shown in **Fig S2**. Data on *dfg10Δ* are from Wilson *et al.* 2024^11^ (**Fig 4D)**. They are displayed here for comparison. (B) Kinetic parameters for Env9 were determined by measuring dolichol formation after incubation of the indicated concentrations of dolichal with 1 mmol/L NADPH and 2mg/mL recombinant Env9 for 5 min at 30°C (means ± SEM; n= 3). (C) Formation of dolichol from dolichal was measured after incubation of 4 μmol/L dolichal with 1 mmol/L NADPH or NADH and 2 mg/mL recombinant Env9 for 1 h at 30°C. None = no protein added to reaction mix. No inc. = reaction mix not incubated prior to metabolite extraction. (D) Kinetic parameters for Env9 were determined by measuring dolichol formation after incubation at 2 μmol/L dolichal with indicated concentrations of NADPH and 2mg/mL recombinant Env9 for 5 min at 30 °C (means ± SEM; n= 3). (E) Formation of polyprenol from polyprenal was measured after incubation of 4 μmol/L polyprenal with 1 mmol/L NADPH or NADH and 2 mg/mL recombinant Env9 for 1 h at 30°C None = no protein added to reaction mix. No inc. = reaction mix not incubated prior to metabolite extraction. (F) AlphaFold model of Env9 with NADPH bound using the “AlphaFill” ^30^ tool indicating the position of p.Arg48, predicted to stabilize binding of NADPH and likely providing its substrate specificity for this cofactor, in turn conveying a preference for catalysis of a reduction reaction with NADPH, instead of an oxidation reaction with NAD^+^.

To test this hypothesis, we produced recombinant Env9 in *E. coli*. Incubation with dolichal in the presence of the cofactor NADPH led to the production of dolichol and a concomitant decrease in dolichal, indicating that this enzyme can indeed act as dolichal reductase (**Fig 2B-C**) with a K_m_ of approximately 1.9 µmol/L for dolichal (**Fig 2B)** and 3.5 µmol/L for NADPH (**Fig 2D)**. In contrast, we did not observe any activity in the presence of NADH (**Fig 2C**), and we did not observe any substantial or consistent interconversion of polyprenol and polyprenal under the same conditions (**Fig 2E)**. This demonstrated that Env9 can only perform one of the two reactions catalyzed by DHRSX (**Reaction 3, Fig 1A**), and that it only uses the cofactor NADP.

The two activities catalyzed by mammalian DHRSX use, alternatively, NAD^+^ for the conversion of polyprenol to polyprenal and NADPH for the conversion of dolichal to dolichol.^11,16^ Cofactor specificity for NADP is often imparted by the presence of positively charged residues interacting in the catalytic pocket with the additional phosphate group present on this nucleotide compared to NAD. The AlphaFold model of Env9 contains an arginine (Arg48) residue that is expected to favor the binding of NADP at the active site. In contrast, the corresponding position is an asparagine residue (Asn75) in *H. sapiens* DHRSX, which could explain DHRSX’s promiscuity for NAD and NADP. (**Fig 2F**).

### Ablation of ENV9 leads to defective N-linked glycosylation

We have previously shown that inactivation of *DHRSX* or *SRD5A3* in human cells and deletion of *DFG10* in yeast lead to defective N-glycosylation due to either accumulation of polyprenol and/or polyprenal, or a decrease in dolichol.^3,11^ To assess the N-glycosylation capacity of *env9Δ* cells, SDS-PAGE mobility of the N-glycosylation marker carboxypeptidase Y (CPY) was assessed. CPY is a yeast protease with four N-glycosylation sites that upon SDS-PAGE analysis shows up to five distinct bands corresponding to its intact fully glycosylated mature form (mCPY) and its hypoglycosylated forms.^31,32^ This has been well characterized in Dfg10-deficient yeast.^3,14^ Env9-deficient yeast exhibited a mild but detectable CPY glycosylation defect in which its hypoglycosylated forms −1 and −2 are clearly visible (**Fig 3A-B**). Reintroduction of *ENV9* restored CPY migration profile to that observed in control cells. The expression of DHRSX, which is able to perform dolichal reductase activity in yeast *env9Δ* cells, also restored CPY glycosylation (**Fig 3A-B**). These data demonstrate that *S. cerevisiae* Env9 dolichal reductase activity is necessary to support sufficient dolichol synthesis for N*-*glycosylation.

**Figure 3.**
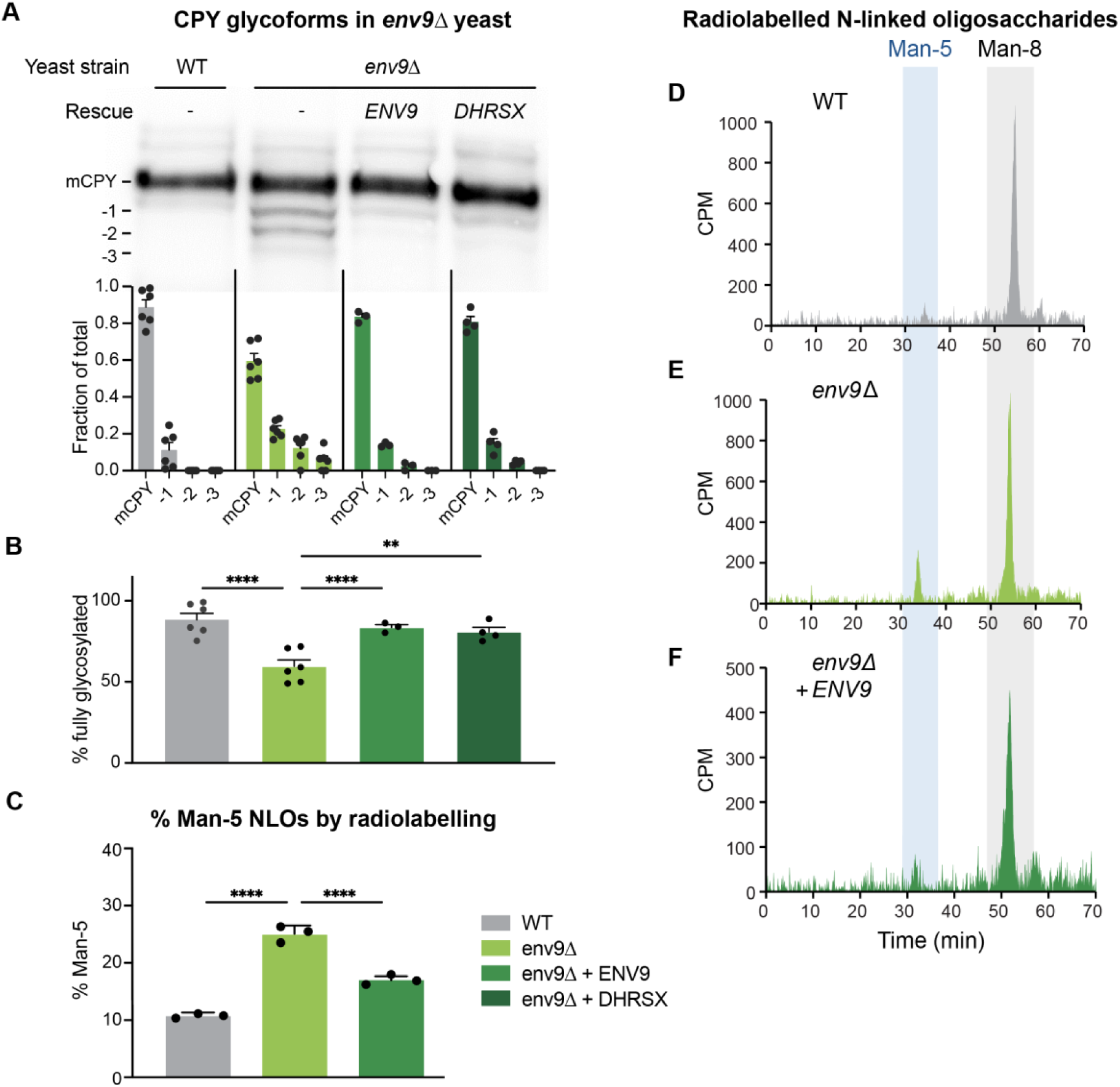
*env9Δ* cells are defective in N-linked glycosylation. (A) Immunoblot analysis of carboxypeptidase Y (CPY) from whole cell protein extracts of wild-type (WT) (BY4741) or *env9Δ S. cerevisiae*, complemented with plasmids carrying *ENV9* or *DHRSX* (indicated as Rescue). Carboxypeptidase Y contains four N-glycosylation sites which are underglycosylated when the N-glycosylation pathway is disrupted. The degree of underglycosylation is indicated by numbers to the left of the immunoblot. Fraction of total indicates the proportion of each band compared to all signal derived from CPY immunoblotting in each lane (means ± SEM; n= 4-6). (B) Relating to **Fig 3A**, the percentage of CPY with four occupied N-glycan sites (means ± SEM; n= 4-6; **p< 0.01; n=3; ****p< 0.0001). (C) Ratios of the abundance of Man-5 to Man-8 N-linked oligosaccharides (NLO) detected in metabolic labeling experiments in wild-type (WT), *env9Δ*, and *env9Δ* yeast cells complemented with *ENV9* (shown in D–F). Means ± SEM, n= 3–6; ****p< 0.0001). CPM = counts per minute. Note that the areas under the curves for Man-5 and Man-8 peaks were normalized according to the number of mannose residues before calculating the ratios. (D) to F. N-linked oligosaccharide (NLO) HPLC profiles obtained from wild-type (D), *env9Δ* (E), *env9Δ* with a plasmid carrying *ENV9* (F) yeast cells labeled with 100 µCi [2-^3^H] mannose, showing a mild accumulation of Man-5 in *env9Δ* cells.

### *env9Δ* cells accumulate abnormal N*-*linked oligosaccharides

When nascent N-linked oligosaccharides (NLOs) are monitored in DHRSX-deficient human cell lines, truncated Man_4-6_GlcNAc_2_ species are observed.^11^ This can be quantified by monitoring the ratio of Man_5_GlcNAc_2_ (Man-5) to Man_9_GlcNAc_2_ (Man-9) NLOs. Analysis is performed by incubating cells with radioactive labelled [2-^3^H]-mannose. The NLOs are extracted, separated by HPLC, and their radioactivity is monitored as they elute. We compared wild-type (WT) control yeast and *env9Δ* cells. In WT yeast, the most abundant NLO identified was Man_8_GlcNAc_2_ (hereafter Man-8) (**Figure 3D**). In *env9Δ* yeast, a slight but significant increase of immature NLO species corresponding to Man_5_GlcNAc_2_ (hereafter Man-5) was observed (**Fig 3E**). Man-5 N-glycans constitute 10.4% of total NLOs in WT yeast, but 24.5% in *env9Δ* yeast (**Fig 3C**). This accumulation of Man-5 species was suppressed by overexpressing *ENV9* (**Fig 3C & 3F**). These observations indicate that immature glycans are transferred onto proteins in *env9Δ* yeast. This is similar to, but much less pronounced than, the defects observed in DHRSX-deficient human cells.^11^

### Genetic interaction network analysis reveals Tda5 and Ydl114w as likely yeast polyprenol dehydrogenase candidates

The identification of Env9 as a dolichal reductase left an obvious question: which enzyme performs the conversion of polyprenol to polyprenal (**Reaction 1**, **Fig 1A**)? Given that mammalian DHRSX also performs this activity^11,16^, a strong candidate would be a yeast oxidoreductase showing both substantial sequence similarity to DHRSX and negative genetic interaction with other enzymes involved in dolichol biosynthesis. To explore this, we further examined the high-throughput genetic interaction data earlier used to identify *ENV9*^23–25^. In this data, *ENV9* showed a strong negative genetic interaction with *TDA*5 (**Fig 1C**), which encodes a member of the short-chain dehydrogenase/reductase (SDR) family that shares sequence similarity with Env9 and DHRSX (**Fig 1B, Table S1**). Moreover, *TDA5* showed negative genetic interactions (**Fig 4A, Table S4, Table S5**) with genes important for dolichol synthesis (*ENV9*, *NUS1*), N-glycosylation (*ALG14, SEC59*), and GPI anchor synthesis (*GPI18*, *GPI16*, and *GPI11*) (**Fig 4A, green text**), as well as *YDL114W*, which encodes another yeast SDR related to Env9 and Tda5 (**Fig 1B, Table S1, Fig 4A**).

**Figure 4.**
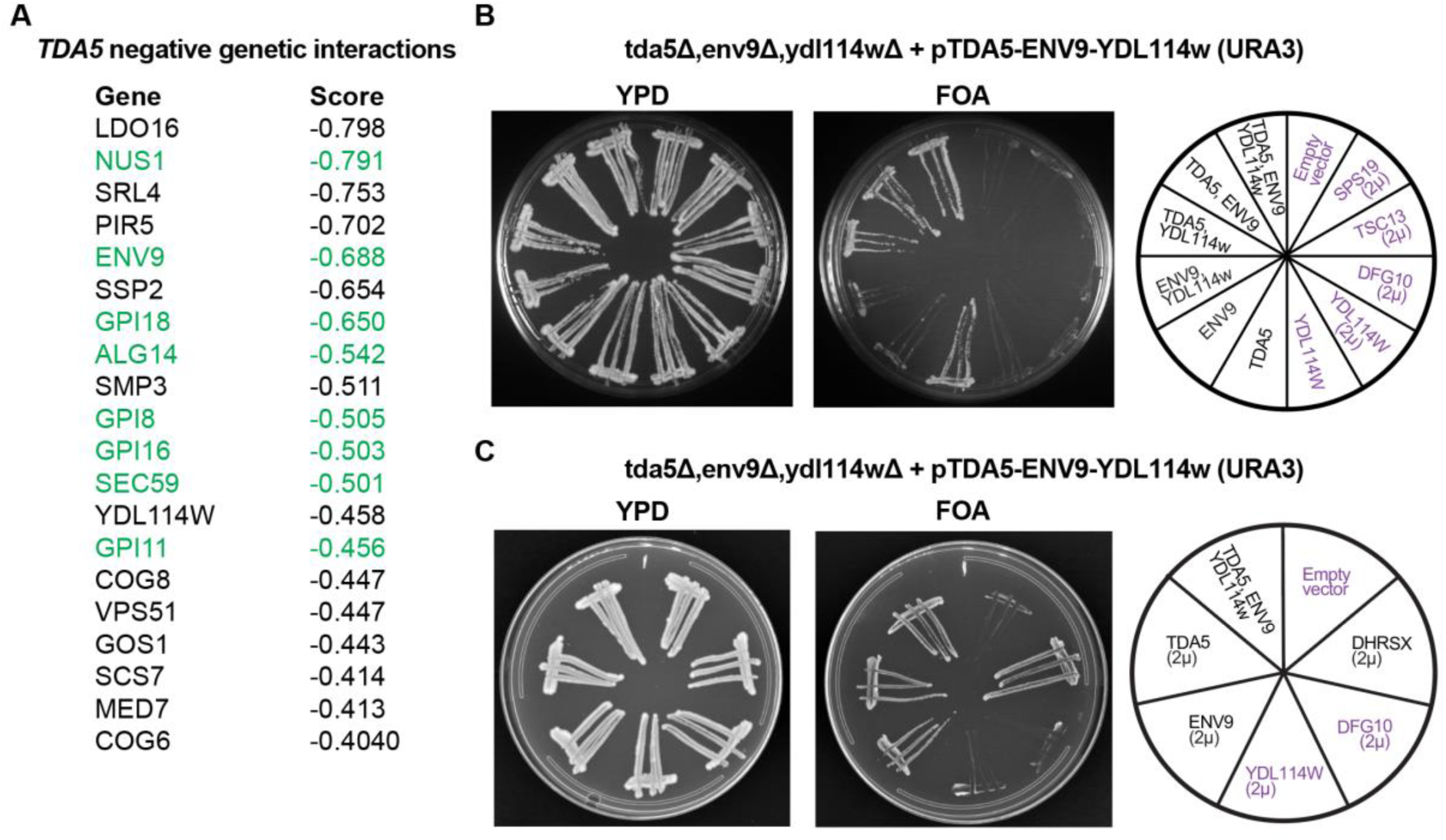
Genetic interaction analysis reveals Tda5 and Ydl114w as likely yeast polyprenol dehydrogenase candidates. (A) Ranked list of 20 strongest negative genetic interactors of *tda5-supp-1*(*sum1*) identified via genome-wide interaction analysis (Costanzo *et al.* 2015^24^). See Table S4 for p-values and systematic gene names. See Table S5 for systematic gene names, mutation alleles and p-values, as well as information for the alternative *tda5-supp-2*(*hst1*) strain. B and C. The *env9Δ ydl114wΔ tda5Δ* triple deletion strain expressing *ENV9*, *YDL114W* and *TDA5* from a *URA3* single-copy plasmid was transformed with single-copy or high-copy (2μ) *LEU2* plasmids containing the genes indicated in the right side of the panel. The cells were streaked onto complete (YPD) plates or synthetic complete medium containing 1% fluoroorotic acid (FOA). The *URA3* gene enables counter selection of cells on FOA. Growth was assessed after 2 days and monitored for 9 days.

These observations were again corroborated by SAFE^24^. In this analysis, we found that areas of cellular function most enriched in negative genetic interactors of *TDA5* included either glycosylation itself, or vesicular trafficking, a process intrinsically linked to glycosylation processes in yeast (**Figure S4A-B**). Overall, these analyses suggested Tda5 might be involved in dolichol biosynthesis and glycosylation, and that Ydl114w might share this function.

### Validating the genetic interactions between *TDA5*, *ENV9* and *YDL114W*

Previous large-scale studies had revealed that TDA5-deficiency rapidly leads to the emergence of loss-of-function suppressor mutations (**Table S4** and **Table S5**)^24,25^, consistent with an important role for viability or proliferation. To explore the functional relationship between *ENV9*, *TDA5* and *YDL114W*, we constructed a yeast strain (KG804) with genomic deletion of all three genes and growth supported by a single vector expressing *ENV9*, *TDA5*, *YDL114W* from their native promoters. This vector carried the counter-selectable *URA3* marker. To interrogate which of these three genes were required for survival and growth, we transformed this strain with another vector that carried different combinations of the three genes. We subsequently selected against the *URA3* plasmid with FOA (**Fig 4C**). In the presence of FOA, no growth was observed for cells carrying the empty vector (EV), demonstrating that the triple knockout strain was not viable. As expected, expression of all three genes together allowed for normal growth. Cells carrying a plasmid that expressed both *ENV9* and *TDA5* also grew normally, indicating that *ydl114wΔ* cells were viable.^33^ Of note, expression of *TDA5* alone, from a single-copy plasmid, was sufficient to maintain growth, whereas *ENV9* expression led to limited growth, and *YDL114W* expression led to no growth. As observed for *TDA5*, human *DHRSX* expression in the triple knockout strain led to normal growth, demonstrating that *DHRSX* can compensate for the loss of both *TDA5* and *ENV9* (**Fig 4D**). The high copy (2μ) expression of *YDL114w* in triple knockout strain led to very limited growth after 7 days of incubation (data not shown). In contrast, high copy expression of the oxidoreductases *TSC13* and S*PS19* did not allow growth. *TSC13*, which encodes a very-long-chain enoyl-CoA reductase, was of interest because it shares structural similarity with *DFG10*. *SPS19*, which encodes a peroxisomal 2,4-dienoyl-CoA reductase, was of interest because it is another yeast SDR with weak similarity to *ENV9* and *TDA5* (**Table S1, Fig 1B**). These observations allowed us to conclude that Tda5 likely displays polyprenol dehydrogenase activity and potentially also some dolichal reductase activity.

### Tda5-deficient *S. cerevisiae* accumulate polyisoprenoids indicative of a polyprenol dehydrogenase deficiency

LC-MS analysis of polyisoprenoids from a *tda5Δ* strain showed a very strong increase of polyprenol levels **(Fig 5A(i))** alongside a depletion of dolichol **(Fig 5A(iv))**. This mirrors the pattern identified in mammalian cells deficient in DHRSX activity. Accordingly, *tda5Δ* yeast accumulated phosphorylated polyprenol **(Fig 5A(iii))**, with a concurrent deficiency of dolichol phosphate **(Fig 5A(vi))**. Expression of yeast *TDA5* or human *DHRSX* lowered polyprenol and polyprenol phosphate levels and increased dolichol and dolichol phosphate levels, close to the levels observed in the empty vector (EV) control cells. These observations were consistent with the hypothesis that Tda5 is the *S. cerevisiae* polyprenol dehydrogenase (**Fig 5B**).

**Figure 5.**
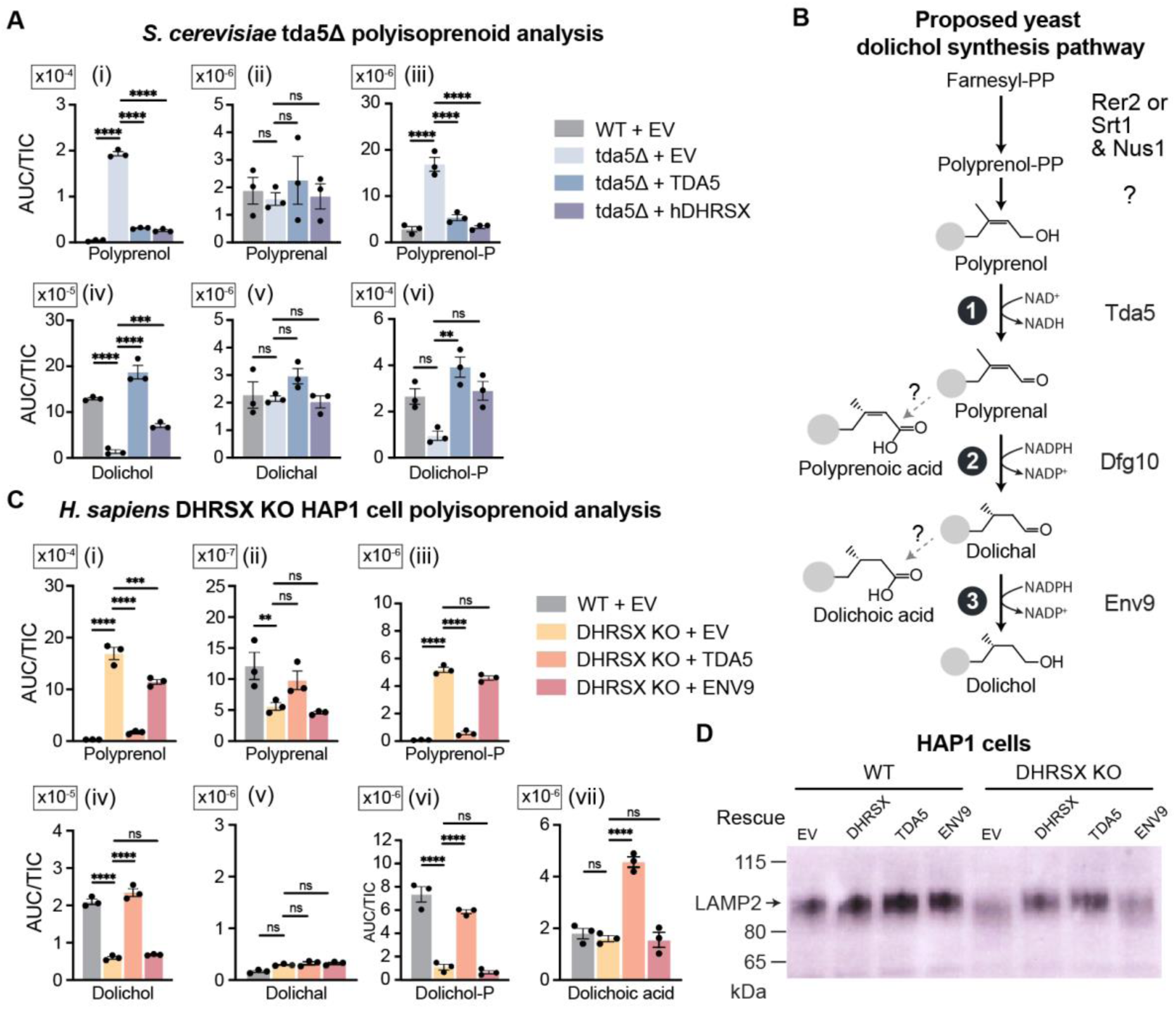
Tda5 is a *S. cerevisiae* polyprenol dehydrogenase. A. Polyisoprenoid species in wild-type and *tda5Δ* and complementations with either TDA5 or hDHRSX. Data represent area under the curve (AUC) normalized to the total ion chromatogram (TIC) (means ± SEM; n=3; **p< 0.01; n=3; ***p< 0.001; ****p< 0.0001). One polyisoprenoid species is shown with 16 isoprene repeats, but additional chain lengths and chromatograms are shown in **Fig S5A**. B. The revised *S. cerevisiae* dolichol synthesis pathway with Tda5 as a polyprenol dehydrogenase, Dfg10 as a polyprenal reductase, and Env9 as a dolichal reductase. C. Polyisoprenoid species in wild-type and DHRSX KO HAP1 cells, and respective complementations. Data represent area under the curve (AUC) normalized to total ion count (TIC) (means ± SEM; n=3; **p< 0.01; n=3; ***p< 0.001; ****p< 0.0001). One polyisoprenoid species is shown with 16 isoprene repeats, but additional chain lengths and chromatograms are shown in **Fig S5B**. D. Immunoblot analysis shows increased LAMP2 mobility indicative of hypoglycosylation in DHRSX KO HAP1 cells. Stable re-expression of WT TDA5 led to a migration of LAMP2 comparable to that in WT HAP1 cells.

To further examine the role of Tda5, we explored its function when expressed in human cells, where DHRSX catalyzes both the polyprenol dehydrogenase and dolichal reductase activities. If Tda5 was specifically a polyprenol dehydrogenase, its overexpression in DHRSX-deficient cells would be expected to lead to a reduction of polyprenol levels and an accumulation of dolichal, since this metabolite could not be further metabolized in the absence of DHRSX. Upon expression of *TDA5* in *DHRSX* KO HAP1 cells, both polyprenol (phosphate) **(Fig 5C(i & iii))** and dolichol (phosphate) **(Fig 5C(iv & vi))** levels were restored to those observed in control cells. Surprisingly, we did not observe a concurrent accumulation of dolichal **(Fig 5C(v))**. At first sight, this suggested that overexpression of *TDA5* could fully replace both the polyprenol dehydrogenase and dolichal reductase activities of DHRSX. However, closer inspection revealed that dolichoic acid levels were increased approximately 2.5-fold in comparison to control cells **(Fig 5C(vii))**. It is likely that dolichal is converted to dolichoic acid in human cells by an unidentified aldehyde dehydrogenase, thereby preventing accumulation of dolichal. A similar, but less efficient, reaction seems to occur in yeast, since the increase of dolichal in *env9Δ* cells was also accompanied by increased dolichoic acid levels (**Fig 2A(vi)**). In summary, normal dolichol levels alongside limited accumulation of dolichal and dolichoic acid in *DHRSX* KO cells overexpressing *TDA5* indicate that Tda5 is able to catalyze at least some dolichal reductase activity. However, this is likely with lower efficiency than DHRSX. An alternative possibility is that human cells contain, besides DHRSX, another, less efficient, dolichal reductase.

### *S. cerevisiae* Tda5 restores underglycosylation of LAMP2 in human *DHRSX* KO HAP1 cells

In human cells deficient in DHRSX, increased mobility of the LAMP2 protein in SDS-PAGE indicated a reduced occupation of N-glycosylation sites and/or incomplete glycans^11^. Expression of *TDA5* in *DHRSX* KO HAP1 cells restored the mobility of LAMP2 to that in control cells, whereas expression of *ENV9* did not (**Figure 5D**). Taken together with the changes in polyisoprenoid levels, our data indicate that Tda5 catalyzes the polyprenol dehydrogenase reaction and, to a weaker extent, the dolichal reductase reaction. In contrast, Env9 only catalyzes a NADPH dependent dolichal reductase reaction. Unfortunately, further enzymological characterization of Tda5 catalytic activity was not possible since we were unable to obtain soluble protein.

### *tda5Δ* cells are defective in N*-*linked glycosylation

We investigated the efficiency of N-glycosylation in *tda5Δ* cells by analyzing the SDS-PAGE mobility of CPY. Consistent with a role of Tda5 in dolichol synthesis, a marked increase in hypoglycosylated CPY glycoforms was observed in *tda5Δ* cells, similar to that observed previously in Dfg10-deficient cells.^3^ This was restored to normal by expression of either yeast *TDA5* or human *DHRSX* (**Fig 6A-B**). In contrast, expression of *YDL114W* was unable to restore N-glycan site occupancy of CPY in *tda5Δ* yeast, providing additional evidence that the gene likely does not contribute to synthesis of the dolichol pool used for protein N-linked glycosylation under normal growth conditions.

**Figure 6.**
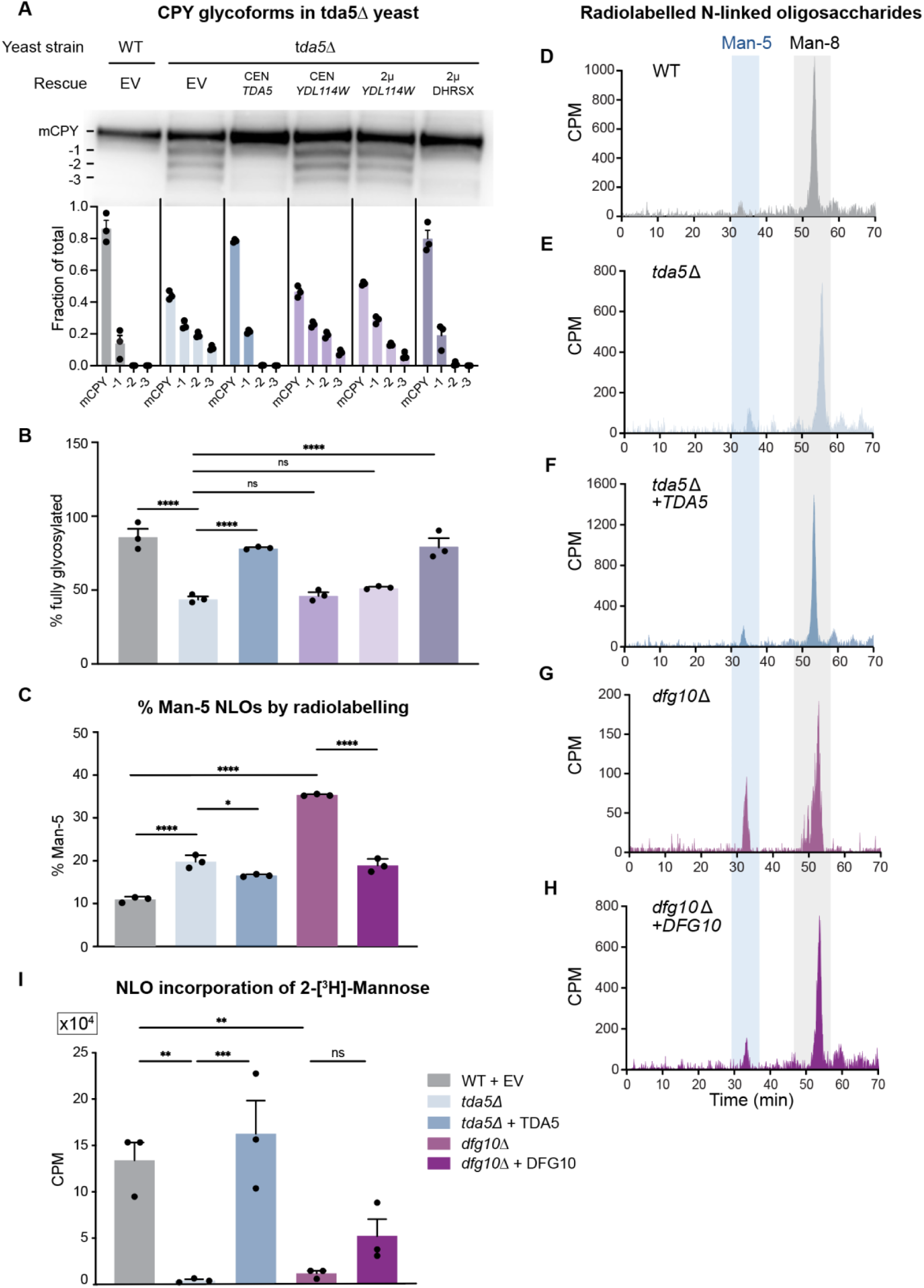
*tda5*Δ cells are defective in N*-*linked glycosylation. A. Immunoblot analysis of carboxypeptidase Y from whole cell protein extracts of WT (BY4741) or *tda5Δ S. cerevisiae*, complemented with plasmids containing *TDA5*, *YDL114W* (in either a CEN or 2µ plasmid) or hDHRSX (indicated as Rescue). Degree of underglycosylation is indicated by numbers to the left of the immunoblot. Fraction of total indicates the proportion of each band compared to all signal derived from CPY immunoblotting in each respective lane (means ± SEM; n= 3). B. Relating to **Fig 6A**, the percentage of CPY with four occupied N-glycan sites (means ± SEM; n= 3; ***p< 0.001; n=3; ****p< 0.0001). C. Relative abundance of Man-5 N-linked to Man-8 N-linked oligosaccharides (NLO) detected in metabolic labeling experiments in wild-type (WT), tda5Δ, and dfg10Δ yeast cells complemented with *TDA5* or *DFG10* (shown in D–H) (2×10^8^ cells were labelled with 100 µCi [2-^3^H] mannose in 50 µl of SC 0.1D medium for 20 min at 25°C). Means ± SEM, n= 3;****p< 0.0001). CPM = counts per minute. Data from WT the same in **Fig 3C** and displayed here for comparison. To ensure accurate quantification, the areas under the Man-5 and Man-8 peaks were normalized to their respective mannose content. D. to H. Representative N-linked oligosaccharide (NLO) HPLC profiles obtained from wild-type (WT), *tda5Δ,* and *dfg10Δ* yeast cells complemented with *TDA5* or *DFG10*, respectively. 2×10^8^ cells were labelled with100 µCi [2-^3^H] mannose in 50 µl of SC 0.1D medium for 20 min at 25°C). Both *tda5Δ* and *dfg10Δ* mutants exhibit an accumulation of truncated NLOs, predominantly Man₅GlcNAc₂. Total radioactivity of purified NLOs was determined before HPLC analysis. Equal amounts of radioactivity was loaded for each sample. E. Total incorporation of radioactivity from [2-3H] mannose into nascent glycoproteins from wild-type (WT), *tda5Δ*, and *dfg10Δ* yeast cells complemented with *TDA5* or *DFG10*. Labelling was performed from 2×10^7^ cells using 10 µCi [2-^3^H] mannose for 10 minutes before extraction of glycoproteins and analysis of radioactivity in CPM (counts per minute) (means ± SEM; n= 3; **p< 0.01; ***p< 0.001; n=3).

### *tda5Δ* cells display a mild accumulation of abnormal N-linked oligosaccharides

We then analyzed newly synthesized N-linked oligosaccharides (NLOs), using metabolic labelling with [2-^3^H]-mannose, identical to the analysis performed for the characterization of Env9 (**Fig 3**). For accurate comparison of NLO profiles, samples were normalized for radioactivity prior to HPLC analysis, with equal amounts of radioactivity injected for each run.

Only a minor accumulation of immature Man-5 NLO was found in *tda5Δ* yeast (19.8% of total NLOs; (**Fig 6C** and **6E**), similar to what we had observed in the *env9Δ* strain (see **Fig 3).** As in WT yeast, the physiological Man-8 structure was the most abundant. In contrast, the Man-5 accumulation was much more pronounced in *dfg10Δ* yeast, accounting for 35.5% of total NLOs (**Fig 6C** and **6G**). This indicated that more immature glycans were transferred onto nascent glycoproteins in Dfg10-deficient yeast compared to Env9- and Tda5-deficient strains. However, the fraction of truncated NLOs was still lower than in DHRSX- or SRD5A3-deficient human HAP1 cells, where the majority of N-linked oligosaccharides are truncated Man_4-6_GlcNAc_2_ species.^11^

### *tda5Δ* and *dfg10Δ* cells exhibit defective mannose incorporation into glycoproteins

Although the quality of NLOs was only mildly abnormal in Tda5-deficient yeast, N-glycan site occupancy of CPY was more severely disrupted. To accurately quantify [2-³H]-mannose incorporation into newly synthesized ER glycoproteins, while minimizing interference from mannan synthesis, the radiolabeling protocol was optimized. Specifically, the incubation period was reduced while the radioactivity concentration per 1×10^6^ cells remained constant. Using this adjusted method, we observed a dramatic reduction in incorporated [2-^3^H]-mannose in both *tda5Δ* and *dfg10Δ*, from 133,776 ±19,526 CPM in control yeast to 4,302 ± 1,285 CPM in *tda5Δ* and 11,887 ± 2,904 CPM in *dfg10Δ* yeast (**Fig 6I**), fully rescued in complemented *tda5Δ* and partially in *dfg10Δ.* These findings indicate that the severe underglycosylation observed in Tda5-deficient strains stems from disrupted dolichol metabolism. This is likely driven by the marked accumulation of polyprenol/ polyprenol-P and the concurrent depletion of dolichol/dolichol-P in *tda5Δ* mutants. Supporting this, prior research has shown that several LLO pathway enzymes exhibit significantly reduced efficiency when utilizing polyprenol, rather than dolichol, as a sugar carrier.^34–37^

### Defective mannan synthesis in yeast dolichol synthesis mutants is driven by both quantity and quality of N-linked oligosaccharides

In *S. cerevisiae*, N-glycans are extended into branched mannans, which are critical structural components of the cell wall.^38,39^ Man_8_GlcNAc_2_ *N*-linked glycans serve as substrates for further elongation in the Golgi to form α-linked polymannosides containing approximately 200 mannose residues. In *S. cerevisiae*, these polymannosides consist of a long backbone of α-1,6-linked D-mannopyranose units, branched with short side chains of α-1,2-linked mannose residues, often further capped by terminal α-1,3-mannose residues.^40^ Mannoproteins constitute 30-50% of the dry mass of the cell wall in yeast^41^, of which up to 90% consists of glycans.^42^. Given this structural importance, we investigated whether N-glycosylation defects in yeast dolichol synthesis mutants could also result in reduced mannan content.

Total mannan levels were quantified across the different yeast mutant strains. In *tda5Δ* cells, which exhibited a reduced quantity (**Fig 6A&I**) but nearly normal quality of NLO (**Fig 6E**), we observed a moderate reduction in total mannan weight compared to control cells (WT = 1.52 ± 0.15 μg/10^6^ cells; *tda5Δ* = 0.97 ± 0.07 μg/10^6^ cells (P = 0.0086)) (**Fig 7A**). In contrast, *dfg10Δ* cells, which exhibited defects in both NLO quantity and quality, showed a marked depletion of mannan (0.31 ± 0.11 μg/10^6^ cells (P = <0.0001 vs. WT)). In *env9Δ* cells, where NLO defects were mild with regards to both quantity and quality, mannan levels remained similar to WT (1.66 ± 0.11 μg/10^6^ cells). Finally, we compared these profiles to *alg3Δ* cells, which exhibit a relatively mild defect in NLO quantity but strongly accumulate truncated, linear Man-5 NLOs.^43^ In these mutants, moderately low mannan levels were identified, similar to those in *tda5Δ* cells (0.84 ± 0.08 μg/10^6^ cells (P = 0.0034 vs. WT)).

**Figure 7.**
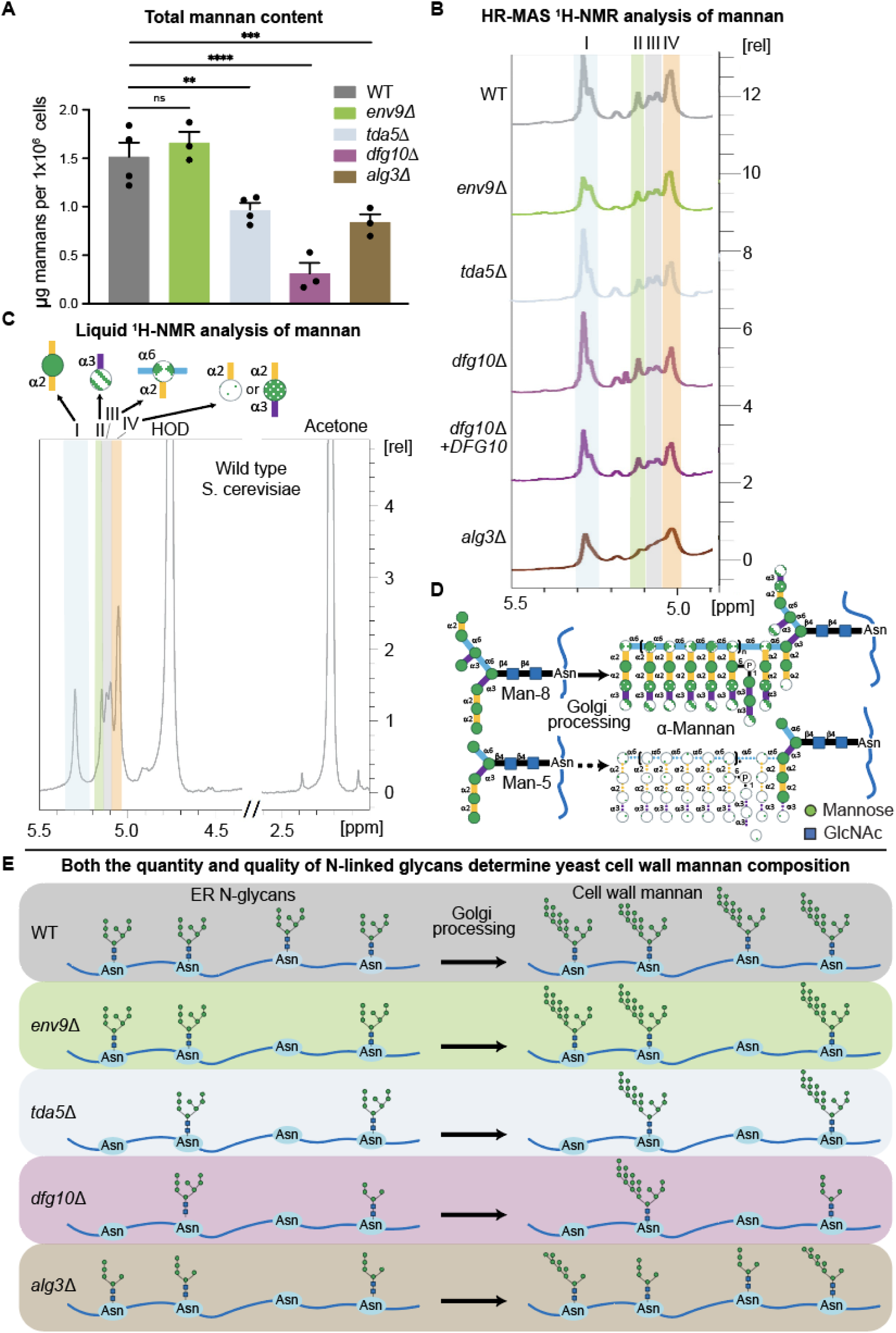
Defective mannan synthesis in yeast dolichol synthesis mutants is driven by defects in both quantity and quality of N-linked oligosaccharides. **A.** Total mannan content across wild-type and dolichol-cycle mutant strains. Weight of extracted mannan (µg/10^6^ cells) for WT, *env9Δ*, *tda5Δ, dfg10Δ* and *alg3Δ*. Means ± SEM, n= 3; * p< 0.05; ***p< 0.001. B. Analysis of yeast dolichol synthesis mutants using High-Resolution Magic Angle Spinning (HR-MAS) NMR (800 MHz). C. Liquid NMR (400 MHz) analysis of mannans extracted from WT and *dfg10Δ* cells. Indicated are the peaks corresponding to (I) internal α1,2-linked mannose residues (δ_H_ 5.3 ppm); (II) terminal α1,3-linked mannose residues (δ_H_ 5.17 ppm); (III) α1,2,6-branched mannose residues (δ_H_ 5.11 ppm); (IV) terminal α1,2-linked mannose residues and α1,2,3-branched mannose residues (δ_H_ 5.05 ppm). D. Och1, which initiates mannan synthesis by attaching the first α-1,6-linked mannose to the α-1,3-arm of the N-glycan core, exhibits low catalytic efficiency when using linear Man₅GlcNAc₂ (Man-5) as substrate.^46^ E. Schematic representation of the hypothesized mechanism by which impaired dolichol synthesis disrupts mannan polymerization and compromises cell wall structure: i) *env9Δ* cells display almost normal mannan due to only mildly reduced quality/mildly defective quantity of NLOs; *tda5Δ* cells display moderately reduced mannan in the cell wall due to a severely reduced quantity but only mildly defective quality of NLOs; *dfg10Δ* yeast display a severe cell wall/mannan synthesis defect as a result of both a severely reduced quantity and abnormal quality of NLOs. *alg3Δ* cells display moderately reduced and abnormally processed (lacking II: α1,3-mannose and III: α1,2-substituted α1,6-mannose residues) mannan due to mildly reduced quantity but severely defective quality of NLOs.^43^

These results indicated that the observed mannan deficiency was dependent not only on the number of fully occupied N-glycosylation sites but possibly also on the quality of NLOs – in this case, the presence of the Man-5 glycoform. To investigate this, we sought to determine whether this mannan deficiency stemmed from an overall decrease in N-linked mannans or abnormal polymannan structures.

To obtain structural insights, we used High-Resolution Magic Angle Spinning (HR-MAS) NMR at 800 MHz to analyze intact yeast cells in the solid phase to obtain structural insights (**Fig 7B**). A primary advantage of HR-MAS NMR is its ability to detect chemical shifts nearly identical to those of liquid-state NMR, while circumventing the need for intensive chemical extraction or purification. By facilitating the direct observation of the cell wall environment *in situ*, this approach preserves the native structural integrity of the glycans and avoids conformational disruptions inherent in traditional processing.^44^ The spectral region between 5.0 and 5.5 ppm displayed a complex profile of signals corresponding to cell wall mannans (see Vinogradov *et al.* 1998^45^). To validate these signals as mannans, we compared the HR-MAS data with Liquid NMR (400 MHz) results from extracted mannans. The profiles for WT and *dfg10Δ* cells acquired using HR-MAS were nearly identical to those acquired using Liquid NMR (**Fig 7C**), confirming that solid phase HR-MAS accurately captures the structural fingerprint of the cell wall *in situ*.

Four characteristic signals were detected in the anomeric proton region (4.8-5.4 ppm) and were designated as signal I, II, III and IV, corresponding to α-mannose residues (**Fig 7B-C**). These signals are four major epitopes: I) internal α1,2-linked mannose residues (δ_H_ 5.3 ppm); II) terminal α1,3-linked mannose residues (δ_H_ 5.17 ppm); III) α1,2,6-branched mannose residues (δ_H_ 5.11 ppm) and; IV) terminal α1,2-linked mannose residues and/or α1,2,3-branched mannose residues (δ_H_ 5.05 ppm). These are illustrated in **Fig 7C**. HR-MAS NMR spectra of WT strain and the mutant strains *env9Δ*, *tda5Δ*, *dfg10Δ*, and d*fg10Δ* + *DFG10* showed no detectable differences compared with the WT (**Fig 7D**), representing the standard branching architecture of mannan in the fungal cell wall. In contrast, *alg3Δ* yeast exhibited a loss of signals II and III, indicating the absence of terminal α3-mannose capping epitope and alterations in the mannan’s backbone. The inability of *alg3Δ* cells to perform normal extension of the mannan backbone is likely due to the prevalence of linear Man-5 NLOs in these yeast.^43^

Our data lead to the conclusion that, among the strains analyzed, *dfg10Δ* yeast display the most severe cell wall defect due to a failure in both the quantity and the quality of NLOs. In contrast, mannan levels are normal in *env9Δ* cells (due to only marginal defects in NLOs quantity and quality) or only mildly reduced in *tda5Δ* and *alg3Δ* cells that show severely reduced NLO quantity and severely defective quality of NLOs, respectively^43^ (see **Fig 7E)**.

## Discussion

### *S. cerevisiae* polyprenol to dolichol conversion uses the same metabolic pathway as *H. sapiens*, but with three enzymes instead of two

Previously, we have shown that Dfg10, the *S. cerevisiae* ortholog of human SRD5A3, is a polyprenal but not a polyprenol reductase.^11^ This indicated that the dolichol synthesis pathway in lower eukaryotes might be arranged similarly to that in mammals. In this work, we confirm that *S. cerevisiae* dolichol synthesis occurs first via the oxidation of polyprenol to polyprenal, then reduction of polyprenal to dolichal, and finally the reduction of resultant dolichal to dolichol. We found that the polyprenol dehydrogenase reaction is almost certainly catalyzed by Tda5, and the dolichal reductase reaction mainly by Env9. These are two oxidoreductases whose natural substrates were previously unidentified. The existence of this three-step detour for dolichol synthesis in both yeast and mammals means that it is likely conserved in all eukaryotes.

### Why two enzymes instead of one?

In the mammalian dolichol synthesis pathway, DHRSX catalyzes two nonconsecutive steps using both NAD^+^ and NADPH as cofactor for different activities. We have shown here that the situation is different in yeast, where two distinct oxidoreductases, Tda5 and Env9, perform polyprenol dehydrogenase and dolichal reductase activities, respectively.

There are two possible reasons that yeast may require two separate enzymes for these activities: Firstly, multicellular organisms maintain quite stable extracellular metabolite levels, allowing individual cells to exist in a constant environment. Thus, intracellular metabolite levels and the redox states of NAD and NADP are more stable than in unicellular organisms, which live in extracellular milieus of highly variable compositions. Additionally, in mammals the NAD^+^/NADH ratio is >500:1^51–53^ in the cytosol, but this ratio is estimated to be lower in *S. cerevisiae* (30-50:1) ^47–49^. Furthermore, it fluctuates widely in yeast and other unicellular organisms in response to environmental factors such as shifts between fermentation and mitochondrial respiration.^18,19^ A hypothetical *S. cerevisiae* polyprenol dehydrogenase able to use both NAD^+^ and NADPH as cofactor would also efficiently catalyze the reverse reduction reaction of polyprenal to polyprenol in situations where the reduced cofactors NADH and NADPH are present at relatively high concentrations. This may impair sufficient dolichol synthesis to support glycosylation pathways.

Secondly, beyond the usage of dolichol for glycosylation, *S. cerevisiae* also uses long chain polyprenol containing 18-24 isoprene units, which is only partially converted into dolichol, during sporulation. These compounds are required for the activation of Chs3, the chitin synthase used for the formation of the chitosan layer, a critical component of the spore wall.^50^ This long-chain polyprenol is produced via an alternative cis-prenyltransferase (Srt1/Nus1 complex) which is specifically localized to lipid droplets associated with the spore wall.^51^ It is possible that the subcellular localizations of Tda5 or Env9 are tailored to ensure that not all polyprenol is converted to dolichol during sporulation. Env9 is associated with lipid droplets in vegetative cells capable of dolichol synthesis, though its localization during sporulation has not been studied.^21^

### Residual dolichol implies redundancy in the dolichol synthesis pathway

Despite the observed N-glycosylation defects in *env9*Δ, *dfg10*Δ, and *tda5*Δ cells there was considerable residual dolichol, ranging from 6% of control levels in *tda5*Δ, to 60% in *env9*Δ. This implies that other enzymes can perform individual steps of polyprenol synthesis, but less efficiently, as indicated by the accumulation of intermediates. In the case of *env9*Δ, it is possible that its dolichal reductase activity can be partially replaced by Tda5, as indicated by the lethality of concomitant inactivation of tda5 and env9, as well as by the rescue of dolichol levels in DHRSX KO human HAP1 cells by expression of TDA5. This may explain the relatively mild CPY N*-*glycosylation defect in *env9Δ*, but at the same time begs the question: why is a mild reduction of dolichol, by only 40% in *env9Δ*, enough to present with a N-glycosylation defect? One possibility is that the residual dolichol is not present in a subcellular localization where it can efficiently be used for N-glycosylation. However, the subcellular distribution of polyisoprenoids has not yet been studied in enough detail for us to reach a firm conclusion. Of note, residual dolichol is also observed in mammalian cells where DHRSX or SRD5A3 have been inactivated. This indicates that during evolution secondary pathways or enzymatic side activities have been maintained that allow limited dolichol synthesis even when the main enzymes are inactive.

### Divergent N-linked glycosylation strategies in conditions of dolichol scarcity: human cells prioritize site occupancy while yeast favor glycan quality

In our previous study, we showed that the majority of nascent N-linked oligosaccharides (NLO) in DHRSX or SRD5A3 KO HAP1 cells were truncated Man_4-6_GlcNAc_2_ species.^11^ The most prevalent of these was a linear Man_5_GlcNAc_2_ ‘Man-5’ NLO. This linear Man-5 lipid-linked oligosaccharide (LLO) is usually processed by the ALG3-catalyzed addition of a sixth mannose residue to the α-1,6-linked mannose, on the luminal side of the ER membrane. In conditions of dolichol deficiency and polyprenol accumulation, such as in DHRSX or SRD5A3 KO, the addition of this sixth mannose is disrupted in a process that is not fully clear, but likely due to lower efficiency of ALG3 when acting on polyprenol-linked mono- and oligosaccharides. This leads to N-glycosylation of glycoproteins with linear Man-5^11,16^, also noted in *alg3*Δ *S. cerevisiae.*^52^

In yeast, a moderate rise in Man-5 NLOs is seen when polyprenol-to-dolichol conversion is disrupted in *env9Δ* and *tda5Δ* cells, though significantly less pronounced than in human cells. This suggests that yeast Alg3 can utilize polyprenol-linked mono/oligosaccharide donor/acceptor substrates, unlike its dolichol-dependent human counterpart. This conclusion is supported by recent *in vitro* work showing that *S. cerevisiae* Alg3 can efficiently use phytol-based donor/acceptor substrates.^53^ Phytol carries a double bond between carbons 2 and 3 of its terminal isoprene unit, analogous to the structure of polyprenol. Man-5 accumulation was more pronounced in *dfg10Δ* cells compared to *env9Δ* or *tda5Δ*. This may stem from a specific effect of accumulation of the reactive metabolite polyprenal, either via specific inhibition of Alg3, or by disrupting the membrane environment required for proper glycan assembly. Alternatively, there could be more redundancy for the activities catalyzed by Env9 and Tda5 than that of Dfg10.

By utilizing polyprenol-linked precursors, yeast can bypass defects in dolichol synthesis. This salvage mechanism would maintain sufficient Man-8 NLO**s** to support mannan assembly and maintain a functional cell wall. This necessity arises because the yeast *cis*-Golgi alpha-1,6-mannosyltransferase, Och1, responsible for initiating outer-chain branching of mannan, is inefficient at utilizing the linear Man_5_GlcNAc_2_ N-glycans.^46^ Consequently, the transfer of linear Man-5 to proteins would significantly be more deleterious to *S. cerevisiae* than to mammalian cells. In humans, Golgi mannosidases (MAN1A1, 1A2, and 1C1) can efficiently process Man-5 intermediates into complex and hybrid N-glycans, thereby maintaining functional proteostasis.^47,55^ Furthermore, Man-5 species in mammals can still participate in the ER-associated quality control and folding of newly synthesized glycoproteins. In contrast, yeast cannot extend these linear precursors into mannans for the cell wall. This reduced tolerance likely exerts a selective pressure that favors the observed substrate ‘permissiveness’ of Alg3 toward polyprenol, minimizing the accumulation of truncated glycans even when polyprenol-to-dolichol conversion is impaired.

Maintaining the structural fidelity of N-glycans transferred to nascent glycoproteins imposes a significant physiological cost on yeast, manifested as marked underglycosylation and the loss of specific N-glycan species, particularly in *dfg10Δ* and *tda5Δ* yeast strains. In these mutants, quantitative metabolic labeling with [2-³H]-mannose revealed a profound reduction in glycoprotein mannose incorporation, in accordance with the severe CPY underglycosylation observed. These results suggest that LLO steady-state levels are severely limited by dolichol availability. In these mutants, the restricted dolichol pool must serve both as the oligosaccharide and monosaccharide donors, creating a competitive bottleneck.

In contrast, human cells appear to place more importance on the transfer of truncated Man-5 species. This is likely achieved via a prioritization of dolichol as LLO anchor, as opposed to Dol-P-sugar donor, in order to prevent systemic underglycosylation. This comes at the cost of an altered quality of nascent N-linked glycans, in the form of Man-5 species, but prevents collapse of the N-glycosylation pathway. This strategy may be driven by the stringent mammalian glycoprotein quality control system, which would target underglycosylated proteins for degradation.

## Acknowledgements

We thank Dr. E. Maes and the PAGés platform (Plateforme d’Analyses des Glycoconjugués, US 41 - UAR 2014 - PLBS, France) for their scientific and technical contributions. We also thank the Institut Chevreul (Université de Lille, Villeneuve d’Ascq) for access to the NMR spectrometer.

Funding: the French National Agency (ENIGMncA project, ANR-21-CE14-0049-01) to F.F.; the CNRS IRP GLYCOCDG project to F.F; the Mizutani Foundation for Glycoscience n°250054 to F.F and n°240097 to G.T.B., an FWO senior postdoctoral fellowship (Project ID: 1289023N to M.P.W.); the Jaeken-Theunissen CDG Fund; KU Leuven Global PhD Partnership (to C.R.A., F.F., and G.M. 3M200250). WELBIO 2019, (to G.T.B.), Fondation Médicale Reine Elisabeth, FNRS equipment grant UN06220F and research credit J.0016.23, ARC UCLouvain ARC17/22-079 and ERC consolidator grant #771704 (all to G.T.B.); FWO-FNRS WEAVE program (G061524N to G.T.B., M.P.W., and G.M.), Fonds Baillet Latour (to G.T.B. and E.V.S.). This work was supported by the Fondation pour la Recherche Médicale, grant number SPF 202409019419 to M.M.

## Materials and Methods

### Yeast strains and culture methods

*Saccharomyces* strains used in this study are listed in **Table S6** and plasmids used in this study are listed in **Table S7**. Yeast cells were cultured in 2% (wt/vol) Bacto peptone and 1% (wt/vol) yeast extract supplemented with 2% glucose (wt/vol) (YPD, yeast peptone dextrose medium). Synthetic complete media were made of 0.67% (wt/vol) yeast nitrogen base and 2% (wt/vol) glucose, supplemented with auxotrophic requirements. For solid media, agar (Difco, Voigt Global Distribution Inc, Lawrence, KS) was added at a 2% (wt/vol) final concentration. Transformations, sporulation of the diploid cells, tetrad dissection and plasmid swapping experiments on 5-fluoroorotic synthetic complete medium were performed by standard yeast genetic methods.

A *Saccharomyces* dfg10 deletion mutant was made using a *Saccharomyces* diploid WT BY4743 strain as a starting point. Briefly, deletion of *DFG10* gene was accomplished using the plasmid pUG27 that carries the his3MX6loxP (*loxP*–*his5*^+^–*loxP)* gene disruption cassette according to previous protocol.^54^ A second set of loxP marker cassettes for Cre-mediated multiple gene knockouts in budding yeast. *Nucleic Acids Res.* 2002 30:e23).

PCR primers used to target the *DFG10 gene* were: AGGTTTATCGGCGGACCGATCTGGCTTAAAACGTAGAATCTACCAGTCCA cagctgaagcttcgtacgc (sense) and TTTACCGCGTCTATATATATATGTATATGAATGCCCTAGTGTGCACATTA gcataggccactagtggatctg (antisense).

The pUG27 dfg10 knock-out construct was transformed into BY4743 yeast cells. Transformants able to grow on medium lacking histidine were isolated, and the correct insertion of the deletion cassette was verified by PCR. The heterozygous mutant *dfg10::*hisMX6/*DFG10* was transformed with pTU-DFG10 plasmid (CEN, URA3). Diploid cells were subjected for sporulation and homozygous deletion strain dfg10 transformed with pTU-DFG10 plasmid was isolated.

To isolate env9 knock out strain resistant to nourseothricin, kanMX4 cassette in the strain in the genome of strain ID 2502 was swapped with natMX4 cassette amplified from pAG25 plasmid.^55^ The cells able to grow on medium supplemented with nourseothricin and sensitive to addition of G414 were isolated. To isolate a ydl114w knock out strain resistant to hygromycin B, a kanMX4 cassette in the genome of the strain ID 3811 was replaced with hphMX3 cassette amplified from pAG34 plasmid.^55^ The cells able to grow on medium supplemented with hygromycin B and sensitive to addition of G414 were isolated. Correct insertion of the deletion cassette was verified by PCR.

KG804 (*env9*Δ, *tda5*Δ*, ydl114w*/pTU-TDA5-ENV9-YDL114w was obtain by sequential mating and sporulation of following yeast strains: tda5, env9 to obtain haploid KG801, and then KG801 and ydl114w to obtain final triple deletion strain.

### Functional complementation assay of *S. cerevisiae* deletion mutants to reveal genetic interaction

To phenotypically analyze genetic interaction between *DFG10* and *ENV9*, yeast double deletion strain dfg10-env9 bearing pTU-DFG10 (URA3 selection) plasmid was co-transformed with single-copy vector pTL (leucine selection) carrying yeast *ENV9* gene under native promoter or empty vector and single-copy vector pTM (methionine selection) carrying yeast *DFG10* gene under native promoter or empty vector.

To phenotypically analyze genetic interaction between *ENV9*, *TDA5* and *YDL114w* yeast triple deletion strain KG804 bearing pTU-TDA5-ENV9-YDL114w (URA3 selection) plasmid was transformed with single-copy vector pTL (leucine selection) carrying combination of yeast *ENV9*, TDA5 and YDL114w genes under native promoters or multi-copy vector pKG-GW1 expressing selected genes from strong constitutive *TDH3* promoter. As negative control yeast strain KG804 was transformed with empty *LEU2* vector.

Transformed yeast cells were streaked onto complete YPD plates or synthetic defined medium containing all amino acids, nucleotide supplements, and 1% (w/v) 5-FOA (Gold-Bio). The plates were incubated for up to 7 days at 30 °C to lose the URA3 vector. The growth of cells was monitored over time to assess phenotypic differences.

### Yeast plasmid Constructions

HiFi DNA Assembly method (NEBuilder®, NEB) was used to construct expression vectors. Yeast genomic DNA was used as template to amplify yeast genes. (see supplementary information for cDNA sequences). The reaction included 0.1 pmol of each fragment, 10 mM DTT, 1 mM ATP, 0.5 µL (10 U) of BsaIHF (NEB), 1 µL of T7-DNA-Ligase (3000 U), and 1X Cutsmart buffer (NEB) in a 20 µL volume. The thermal cycling program was: 3 min at 37°C, 35 cycles of 2 min at 37°C and 3 min at 25°C, 5 min at 50°C, and 5 min at 80°C. Five microliters of the reaction were transformed into NEB 5α competent cells, and colonies on selective plates were screened. Plasmid purification used the Macherey-Nagel Nucleospin kit. DNA concentrations were quantified with a nanospectrophotometer ND 1000 (NanoDrop). Constructs were verified by restriction digestion and agarose gel electrophoresis in 0.8% TBE buffer with GelRed (Merck) staining. Construct sequences were confirmed in Sanger sequencing (Eurofins Genomics).

### Isoprenoid species and preparation of dolichal

Dolichol (# 9002000), polyprenol (# 9002100) and polyprenal (# 9002200) were purchased from Avanti Polar Lipid. Dolichal was synthesized by oxidizing dolichol in the presence of pyridinium dichromate (# 214698-100G, Merck life science BV, (Corey and Schmidt, 1979^56^): 300 mL of 1 mg/mL dolichol in chloroform was dried down and resuspended in 200 mL of dichloromethane containing 10 mg/mL of pyridinium dichromate. After shaking at 1000 rpm for 15 min, the preparation was incubated overnight at 20°C. The mixture was centrifuged, and the supernatant was transferred to a new tube. Several cycles of back-extractions were performed by using 200 mL methanol/water (5:3) (MS-grade, Biosolve) to remove pyridinium dichromate until the preparation became translucent. The quality of dolichal synthesis was assessed by LC-MS, showing a high yield of conversion (>95%). Dolichal was stored at −80°C for future enzymatic assays. All isoprenoid solutions represent mixtures with 13–21 isoprene units (with 18 being the most abundant species). Molar concentrations were calculated based on the distribution of chain lengths. Thus, a 5 μg/mL solution corresponds to approximately 4 μmol/L.

### Sample preparation and extraction of metabolites from yeast

*S. cerevisiae* strains were cultured overnight in 20 mL SC-Leu medium at 30°C and 200 rpm for each strain. OD600 was measured to obtain approximately equal amounts of yeast in each sample. Depending on the strain, 9-17 mL of cultures were then centrifuged at 13200 rpm at 4°C. The pellet was immediately plunged into liquid nitrogen, resuspended in ice-cold methanol/water/chloroform (5:3:10) and vigorously homogenized. The homogenate was moved to a 2 mL tube and subjected to 2 cycles of freezing-thawing in liquid nitrogen. The lysate was vigorously shaken for 20 min at 2000 rpm at 4°C, and centrifuged at 13200 rpm for 30 min at 4°C. The lower organic layer was dried down under a gentle stream of nitrogen and resuspended in 100 μL of methanol/isopropanol (1:1) for LC-MS analysis.

### Dimethylation of isoprenoid phosphates using trimethylsilyl diazomethane (TMSD)

The dimethylation of dolichol phosphate and polyprenol phosphate was performed as previously described^57^. The organic fraction containing isoprenoid species was completely dried and resuspended in 200 μL of dichloromethane:methanol (6.5:5.2, v/v). 10 μL of 2 mol/L TMSD (in hexane) was added and incubated for 40 min at room temperature. The incubation was halted by adding 1 μL of acetic acid. Samples were then dried and dissolved in 50 µL methanol:isopropanol 1:1 (V/V) for LC-MS analysis.

### Overexpression and Ni-NTA agarose purification of *S. cerevisiae* Env9 in *E. coli*

pET15b plasmids containing cDNA corresponding to *S. cerevisiae* ENV9 with a 6 x His tag at the N-terminus were transformed into One Shot TOP10 Chemically Competent *E. coli* (Life technologies, ref: C404003). Clones were selected and grown to prepare minipreps. Vectors were then transformed into BL21(DE3) *E. coli.* Colonies were then selected for carbenicillin resistance and grown in 10mL LB medium (100 μg/mL carbenicillin) overnight at 37°C before dilution in 250mL LB medium (100 μg/mL carbenicillin) and further growth for 3-6 hours until an OD600 of 0.6 was reached. Isopropylthio-β-galactoside (IPTG) was then added to a final concentration of 1 mmol/L and the culture was grown for a further 5 h at 37°C before centrifugation at 4000 x g for 20 min to harvest cells. Native purification of 6 x His-tagged Env9 was performed using the Ni-NTA fast start kit (Qiagen ref. 30600) according to manufacturer’s instructions.

### Env9 activity using polyprenal and dolichal

To identify the substrate and the cofactor of ENV9, 4 µmol/L of dolichal or polyprenal were used in the presence of 1 mM NAD(P)H in the aim to measure the formation of their reduced forms; dolichol and polyprenol, respectively. The incubation buffer contained sodium phosphate 20 mmol/L, pH 6.5, Triton X-100 0.2%, phosphatidylcholine 1%, phosphatidylethanolamine 0.2%, glycerol 10%, ß-mercaptoethanol 0.5 mmol/L, KCl 10 mmol/L. The reaction was started by adding the enzymatic preparation at 2 mg/mL, incubated for 1 hour at 30°C and 400 rpm. The assays were stopped by adding methanol/water (5:3). Samples were analyzed by LC-MS after methanol-chloroform extraction. For the determination of *K_M_* of dolichal, the concentrations of 0, 0.2, 0.4, 0.8, 1.6 and 4 µmol/L were used with 1 mmol/L of NADPH. The incubation time was 30 min and temperature 30°C. Concentrations of 0, 5, 20, 100, and 1000 µmol/L of NADPH were used to identify its *K_M_* in the presence of 2 µmol/L dolichal.

### Sample preparation and extraction of metabolites from enzymatic assays

After the end of each respective incubation, 450 μL of ice-cold methanol/water (5:3) was added into the tube followed by 550 μL of ice-cold chloroform. The mixture was vigorously mixed at 2000 rpm for 20 min and at 4°C, followed by a centrifugation at 13200 rpm for 10 min at 4°C. The lower layer containing hydrophobic metabolites was dried down under a gentle stream of nitrogen and resuspended in 50 μL methanol/isopropanol (1:1) before LC-MS analysis.

#### LC-MS analysis of metabolites

LC-MS analysis of organic fractions obtained from cells (or enzymatic assays) was carried out using a method previously described in Kentache et *al*., 2024^16^ and Wilson et *al*., 2024^11^. Briefly, 5 μL of sample was injected and subjected to reverse phase chromatography with an Accucore C30 150 × 2.1 mm column (ref. 27826–152130, ThermoFisher), operated at 45°C on an Agilent 1290 HPLC system. The flow rate was constant at 0.2 mL/min using mobile phase A (60% acetonitrile, and 40% water, 10 mmol/L ammonium formate, and 0.1% formic acid) and B (90% isopropanol, 10% acetonitrile, 10 mmol/L ammonium formate, and 0.1% formic acid (Biosolve)). An Agilent 6546 ion funnel mass spectrometer was used in the positive or negative ionization modes with an electrospray ionization (ESI) (voltage 3500 V, Nozzle voltage 1000V, sheath gas 350 °C at 11 L/min, nebulizer pressure 35 psi and drying gas 300 °C at 8 L/min). Starting from 5 min onwards, one spectrum encompassing a range of 69–1700 m/z was acquired per second, generated from 10772 transients. The mass spectrometer was operated in positive mode for the detection of dolichal, polyprenal, dolichol, polyprenol, dimethylated dolichol-P, and dimethylated polyprenol-P. For the elution, the solvent gradient was: 0–5 min at 90% B; 5–33 min from 90 to 97% B; 33–34 min from 97 to 99% B; 34–35 min from 99 to 90% B. The negative mode was used to detect and measure polyprenoic acid, dolichol-P-hexose, and polyprenol-P-hexose. The elution gradient consisted of: 0–3 min at 30% B; 3–8 min from 30 to 43% B; 8–9 min from 43 to 50% B; 9–18 min from 50 to 90% B; 18–26 min from 90 to 99% B; 26–30 min at 99% B; 30–30.1 min from 99 to 30% B; 30.1–35 min at 30% B. The theoretical m/z values of [M + NH4+] and [M - H+] ions can be found in **Table S1** of Wilson *et al,* 2024^11^. The resulting data were analyzed and processed by the software Agilent MassHunter Qualitative Analysis 10.0 for the identification and the visualization of peaks/metabolites. The quantification was performed by Agilent MassHunter Quantitative Analysis software (Agilent Technologies, CA, USA).

### Recombinant lentivirus production and infection of HAP1 cells

Recombinant lentiviruses were produced in HEK293T cells as previously described^58^ using the vectors pUB81 TDA5. pUB81 is a lentiviral expression vector based on the pLVX-PURO (Clontech) plasmid containing EF-1α promoter. Sequences were verified by Sanger sequencing. Viruses were then used to infect control and DHRSX KO HAP1 cells. HAP1 cells were grown in 6 well plates to reach 60% confluence on the day of infection in IMDM. The medium was replaced by IMDM containing diluted recombinant lentiviruses (lentivirus:IMDM 1:4 V/V) in the presence of 4 μg/mL polybrene. After 24 h of growth the medium was replaced by virus free IMDM containing 2 μg/mL puromycin. For isoprenoid measures by LCMS, cells are seeded in 6 cm plates and cultured for 1 or 2 days until reaching ±90% of confluency. Then, the medium was removed, plates were rapidly washed with ice-cold water and plunged in liquid nitrogen to quench metabolic activity. The frozen dishes were placed on dry ice, and 500 μL ice-cold methanol was immediately added, followed by 300 μL of ice-cold water. The cells were scraped and collected in 2 mL tubes containing 1 mL of chloroform (MS-grade, Biosolve). The 2 mL tubes containing lysates were vigorously mixed at 2000 rpm for 20 min at 4°C, followed by a centrifugation at 13200 rpm for 30 min at 4°C. The lower layer, containing hydrophobic metabolites, was dried down under a gentle stream of nitrogen and resuspended in 60 μL of methanol/isopropanol (1:1) before LC-MS analysis.

### Immunoblotting of protein lysates from yeast

Protein lysates from yeast cells were prepared as described previously.^32^ In brief, RIPA buffer (10 mmol/L Tris-HCl [pH 7.4], 150 mmol/L NaCl, 0.5% sodium deoxycholate, 0.1% SDS, and 1X cOmplete protease inhibitor cocktail (Sigma Aldrich)) was first added at 4°C. The yeast cells were disrupted using 0.5 mm glass beads through five cycles of 30-second vortexing followed by 1-minute cooling on ice. Lysates were collected by centrifugation at 4°C (15,000 rcf for 5 minutes). The total protein concentration was measured with the Pierce BCA protein assay kit (Thermo Fisher Scientific). Proteins were separated by SDS-PAGE and transferred to a nitrocellulose membrane (Invitrogen, LC2000). The membrane was blocked with 5% milk (98022, Sigma-Aldrich) and incubated with the primary antibody against CPY (1:1000; Life technologies A6428) and the secondary Goat-anti-Mouse antibody (P0447, DAKO). The membranes were washed with 1X Tris-buffered saline containing 0.1% Tween-20 (P1379, Sigma-Aldrich). Signal detection was carried out using chemiluminescence on an Amersham ImageQuant 800 imager (Cytiva).

### Immunoblotting of protein lysates from human HAP1 cells

Protein lysates from human HAP1 cells were prepared as described previously^11^ and as performed for yeast cells, with the exception of the mechanical lysis step which was performed by passing the lysates 5 times through a sterile 27G needle prior to centrifugation. Immunoblotting was perform with an antibody recognising LAMP2 (H4B4; sc-18822; Santa Cruz Biotechnology).

### Yeast metabolic labelling

*S. cerevisiae* strains were seeded at an initial OD_600_ of 0.2 in 20 ml of SC medium from a pre-cultured of 5 ml and then grown until mid-log phase (final OD_600_ of 0.6-0.8). The cells were then pelleted by centrifugation at 500 x *g* for 5 min at room temperature (RT), washed two times in 1 mL of SC medium containing only 0.1 % of glucose (SC 0.1D) and then labeled with [2-3H] mannose (American Radiobiolabeled Chemicals, Inc (ARC), Saint-Louis, USO/Isobio, Fleurus, Belgium). For the qualitative analysis, 2×10^8^ cells were pelleted and labeled with 100 µCi of [2-3H] mannose in 50 µl of SC 0.1D medium for 20 min at 25°C. For the quantitative analysis, labeling was performed from 2×10^7^ cells with 10 µCi of [2-3H] mannose in 5 µL SC 0.1D, for 10 min at 25°C.

After metabolic labeling, yeasts were pelleted at 3000 x *g*, 2 min at RT and then washed four times in distilled water. The cells were resuspended in 500 µL lysis buffer (0.1 M tris HCl pH 7.4 containing 4 mM MgCl_2_) and homogenized by glass-bead agitation using 5 disruption cycles (30 sec agitation; 1 min on ice). Lysates were collected by centrifugation (3000 x *g*, 5 min, RT) in a glass tube and then 1 ml of methanol and 1.5 ml of chloroform were successively added to obtain a mixture Chloroform/Methanol/Water (3:2:1 (v/v)). The qualitative analysis of oligosaccharides was performed as described by Wilson *et al*., 2024. For the quantitative analysis, after sequential extraction, the pellet of glycoproteins was transferred directly to scintillation vials, and the total radioactivity was counted by adding 2 mL of Ultima-Gold scintillation liquid (Revvity) using the HIDEX 300 SL counter. The results were expressed in counts per minute (CPM). All experiments were repeated at least 3 times.

### Yeast Mannan Extraction

Yeast cells were cultured in SC medium (*n*=3), harvested at OD600 ≈ 0.8 by centrifugation, and pellets were resuspended in 0.2 M citrate buffer (pH 7.0). Cells were autoclaved at 120 °C for 1 h, centrifuged, and the supernatants pooled after a second extraction cycle. Combined extracts were incubated overnight at 4 °C with an equal volume of Fehling’s solution, and the resulting precipitate was collected, dissolved in 3N HCl, and reprecipitated with ethanol. The mannan pellet was resuspended in water and dialyzed overnight (6–8 kDa, 4 °C) against 500 mL of distilled water before storage at −20 °C and lyophilization to obtain the dry weight.

### Liquid state NMR analysis

Mannan was dissolved in D_2_O (99.96% ^2^H, Eurisotop®) at a concentration of 1mg/mL, and then transferred into 5 mm Shigemi tubes (Allision Park, USA). NMR spectrum was recorded at 298 K at proton resonance frequency of 400 MHz on Avance II Bruker spectrometer equipped with TBI 5mm probe. Proton chemical shifts were referenced to acetone (δ 1H 2.225 ppm), which was used as internal standard. Spectra were processed using Topspin 4.5.0 (Bruker GmbH, Germany).

### HR-MAS analysis

Yeast cells were cultured in SC medium, harvested at OD600 ≈ 0.8 by centrifugation and were washed twice with 1 mL of D_2_O (Eurisotop, St Aubin, France). The cell pellet was resuspended in 50 mL of D_2_O and introduced into a disposable HR-MAS rotor. The rotor was inserted into a 5-mm HR-MAS probe and spun at 8000 Hz on an 18.8 T spectrometer, corresponding to a proton resonance frequency of 800.13 MHz. The spectrometer was equipped with an HCP-type HR-MAS probe, and ^1^H NMR spectra were acquired at 298 K. Pulse programs used for data acquisition were taken from the Bruker pulse program library, with delays and power levels empirically optimized for each experiment. Spectra were processed using Topspin 4.5.0 (Bruker GmbH, Germany).

### Pairwise and multiple protein alignments

Initial identification of *S. cerevisiae* proteins homologous to *H. sapiens* DHRSX was performed using BlastP. Protein sequences were then aligned and visualized using the MAFTT algorithm in Unipro Ugene.^59^ Subsequently, pairwise alignment was analyzed to determine sequence similarity (including gaps) and identify conserved regions.

### Statistical analysis

Analyses were performed using GraphPad Prism version 10.2.3 for Mac (GraphPad Software, La Jolla, CA). Statistical analysis was performed using a Student’s T-test or a one-way ANOVA followed by Tukey’s or Dunnett’s multiple comparisons test.

## Supplementary figures

**Figure S1:**
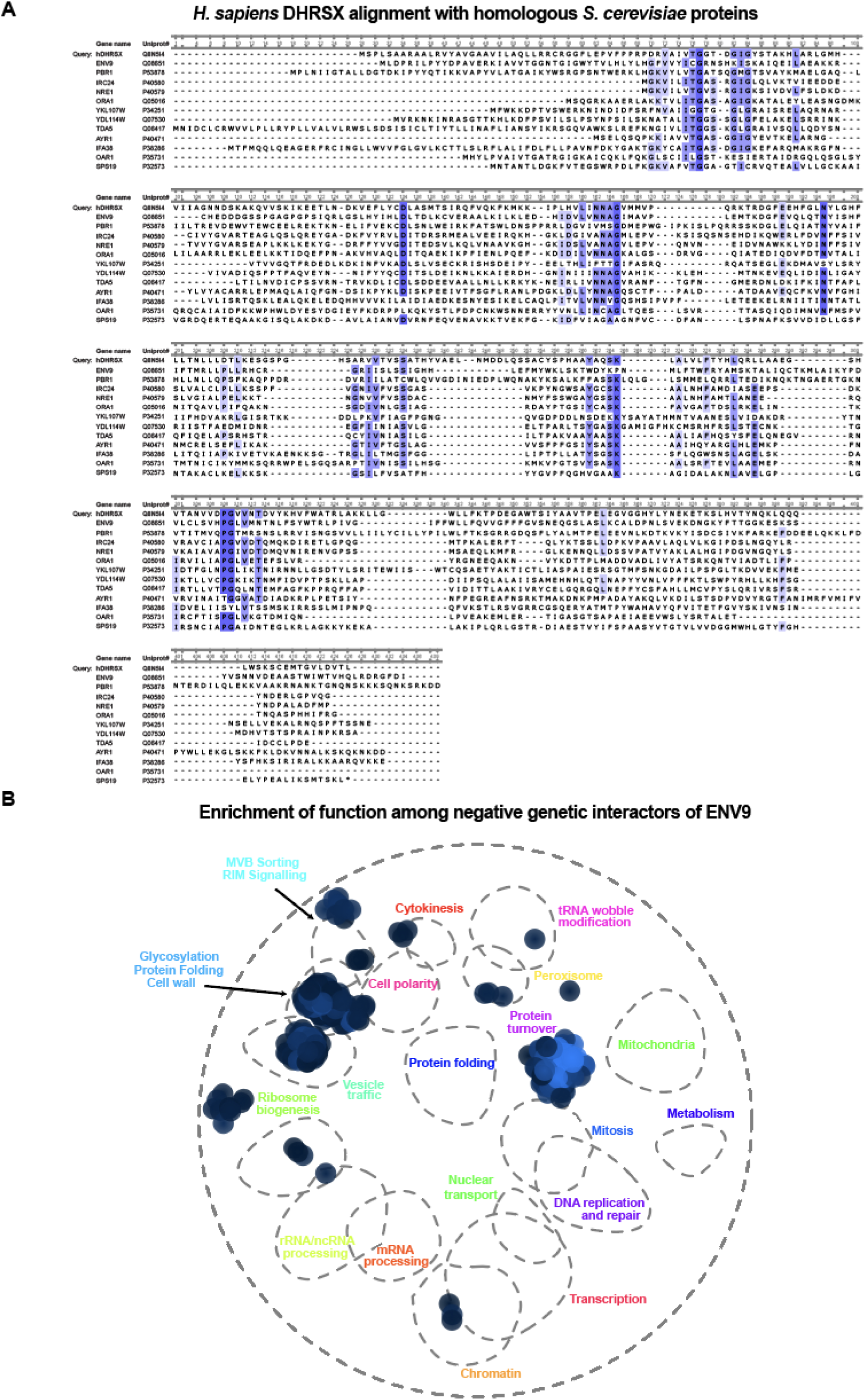
**A:** ClustalW alignment of yeast polyprenol dehydrogenase and dolichal reductase candidates. **B:** ENV9 genetic interaction hotspots acquired by SAFE analysis (Costanzo *et al* 2015) of an env9Δ strain.

**Figure S2:**
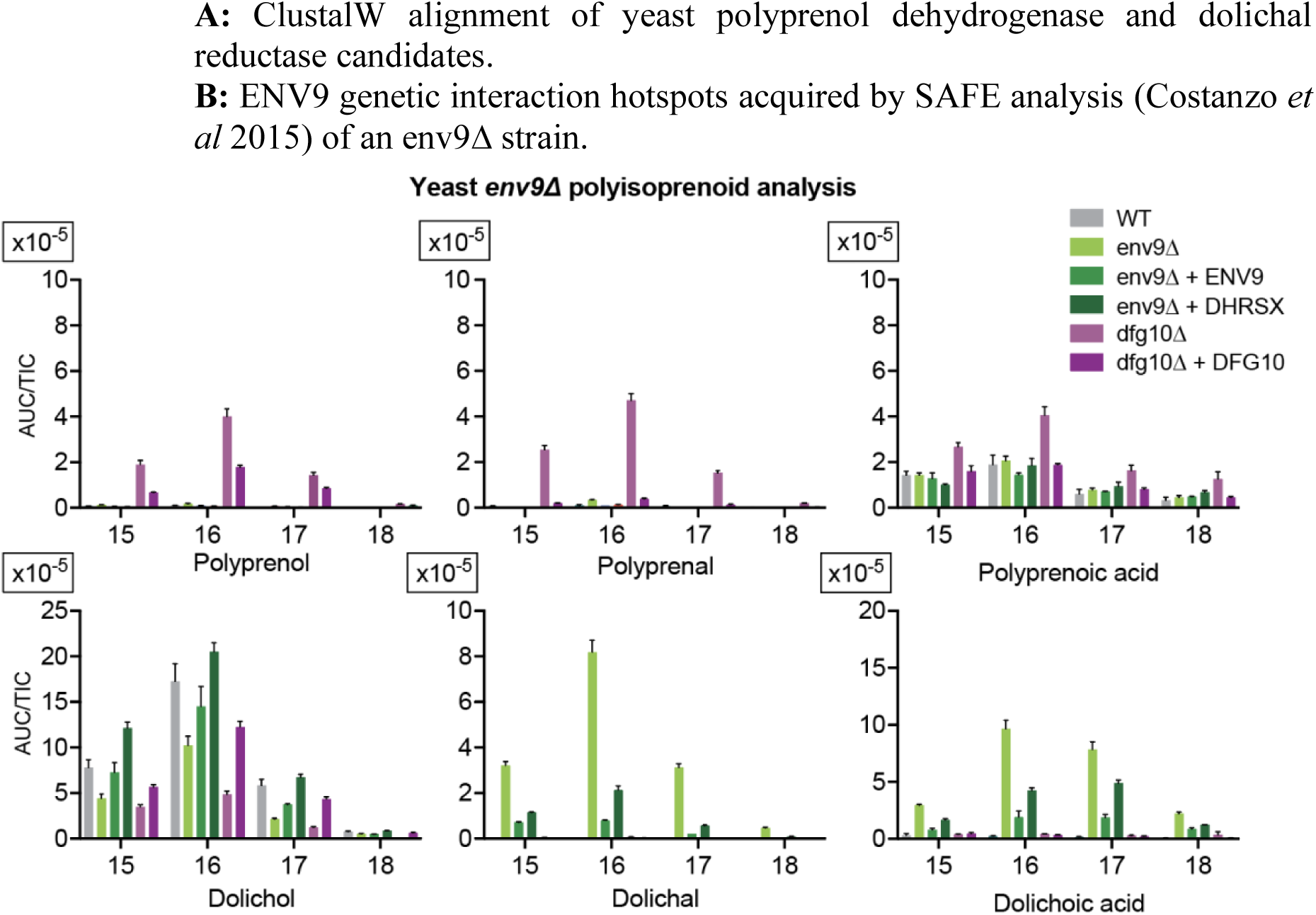
Polyisoprenoid species with 18-21 isoprenyl units in wild-type, env9Δ, dfg10Δ yeast cells and their respective complementations. Data are TIC-normalized AUC (mean ± SEM, n=3). Data on *dfg10Δ* are taken from Wilson *et al.* 2024^11^ Fig S4 and are displayed here for comparison.

**Figure S3:**
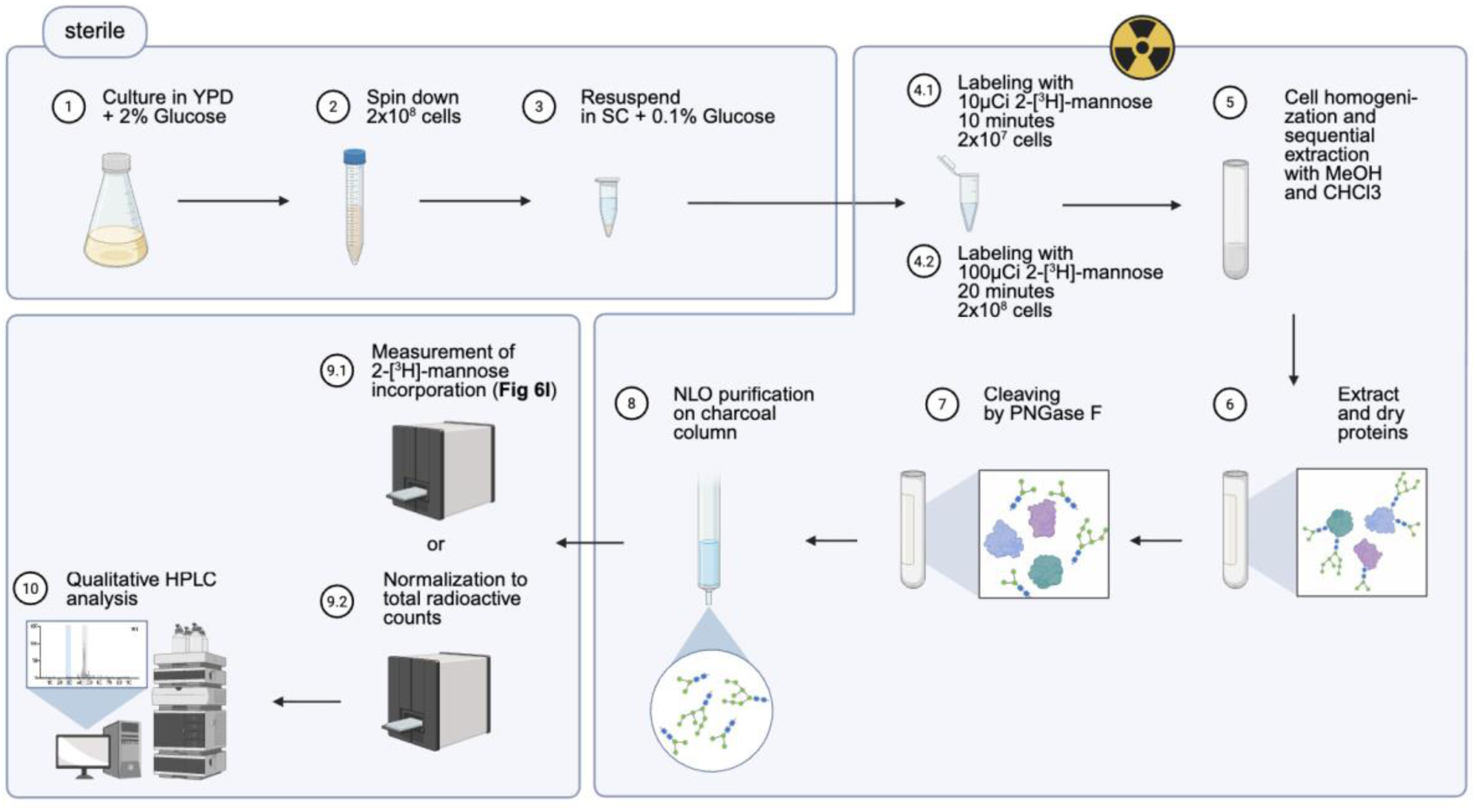
Radiolabelling protocol used for qualititative N-linked oligosaccharide analysis and measurement of 2-[^3^H]-mannose incorporation (Fig. 6I)

**Figure S4:**
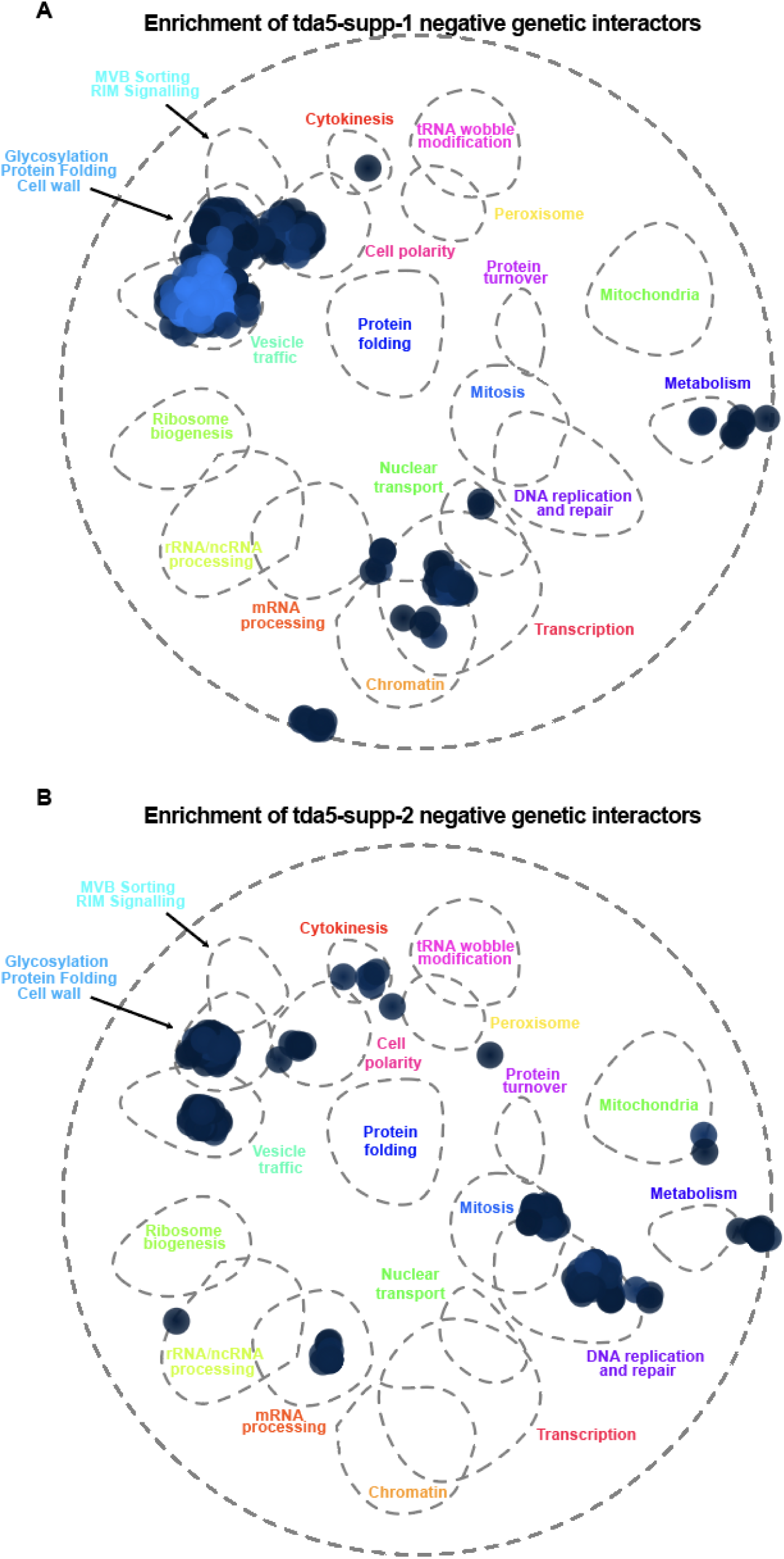
A: tda5-supp-1 genetic interaction hotspots acquired by SAFE analysis (Costanzo *et al* 2015).^24^ B: tda5-supp-2 genetic interaction hotspots acquired by SAFE analysis (Costanzo *et al* 2015).^24^

**Figure S5:**
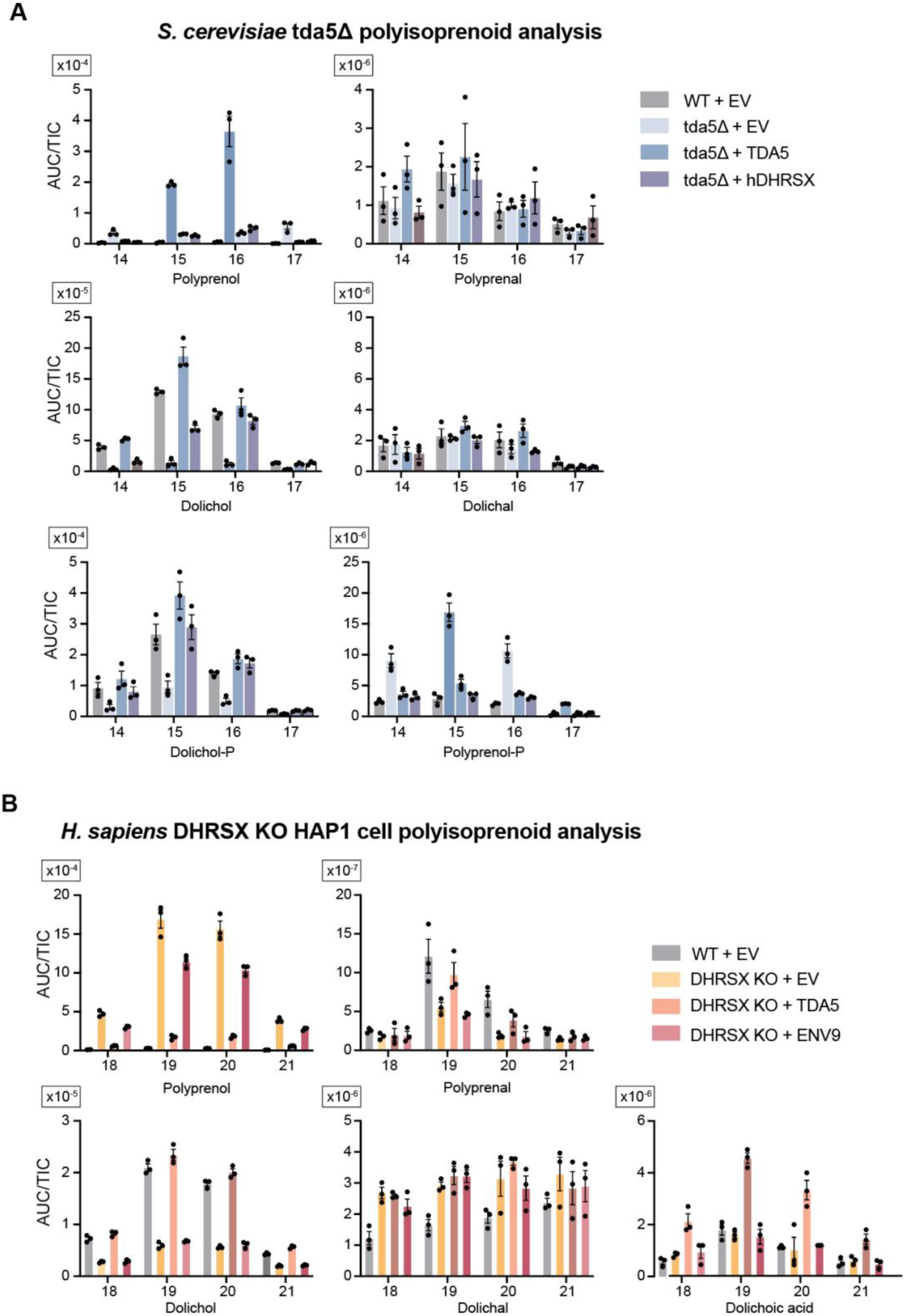
**A:** Polyisoprenoid species with 14-17 isoprenyl units in wild-type and tda5Δ yeast cells and their respective complementations. Data are TIC-normalized AUC (mean ± SEM, n=3). **B:** Isoprenoid species with 18-21 isoprenyl units in control and DHRSX KO, HAP1 cells and their respective complementations. Data are TIC-normalized AUC (mean ± SEM, n=3).

