## Supplementary information for "Dolichol biosynthesis in yeast traces the expanded reaction pathway of higher eukaryotes"

**Table S1: BLASTP analysis of the *S. cerevisiae* proteins most homologous to *H. sapiens* DHRSX**

| **Gene name** | **Systematic gene name** | **Uniprot accession** | **Annotated oxidoreductase activity** | **Identities(%)** | **E-value** |
| --- | --- | --- | --- | --- | --- |
| *ENV9* | YOR246C | Q08651 | Unknown | 27.9 | 7.2e-28 |
| *PBR1* | YNL181W | P53878 | Unknown | 27.8 | 2.8e-19 |
| *IRC24* | YIR036C | P40580 | Benzil oxidoreductase  (NRE1 paralog) | 25.7 | 3.0e-11 |
| *NRE1* | YIR035C | P40579 | Benzil oxidoreductase  (IRC24 paralogue) | 26.6 | 3.4e-09 |
| *ORA1* | YMR226C | Q05016 | 3-hydroxy acid dehydrogenase: catalyzes the reduction of (S)-α-acetolactate to 2,3-dimethylglycerate | 22.4 | 5.4e-08 |
| *-* | *YKL107W* | P34251 | Involved in the detoxification of acetaldehyde, glycolaldehyde, furfural, formaldehyde and propionaldehyde | 19.9 | 1.2e-07 |
| *-* | *YDL114W* | Q07530 | Unknown | 21.9 | 6.7e-07 |
| *TDA5* | YLR426W | Q06417 | Unknown | 23.9 | 2.2e-06 |
| *AYR1* | YIL124W | P40471 | 1-acyldihydroxyacetone phosphate reductase | 28.5 | 3.6e-05 |
| *IFA38* | YBR159W | P38286 | Very-long-chain 3-oxoacyl-CoA reductase | 24.8 | 9.7e-05 |
| *OAR1* | YKL055C | P35731 | 3-oxoacyl-[acyl-carrier-protein] reductase | 23.3 | 0.006 |
| *SPS19* | YNL202W | P32573 | Peroxisomal 2,4-dienoyl-CoA reductase | 21.0 | 0.004 |

**Table S2: Top negative genetic interaction with *ENV9*. Related to Fig 1C.**

| **Gene name** | **Systematic gene name** | **Mutation allele** | **Gene interaction score** | **p-value** |
| --- | --- | --- | --- | --- |
| PRE1 | YER012W | pre1-1 | -0.790 | 5.42e-97 |
| TDA5 | YLR426W | tda5Δ-supp2 | -0.770 | 6.39e-62 |
| RER2 | YBR002C | rer2-5008 | -0.750 | 0.00e+00 |
| RPN6 | YDL097C | rpn6-1 | -0.745 | 4.01e-65 |
| TDA5 | YLR426W | tda5Δ-supp1 | -0.688 | 5.49e-44 |
| PRE7 | YBL041W | pre7-5001 | -0.687 | 0.00e+00 |
| TIM22 | YDL217C | tim22-19 | -0.590 | 2.80e-30 |
| SCL1 | YGL011C | scl1-5001 | -0.588 | 1.99e-46 |
| PRE9 | YGR135W | pre9Δ | -0.543 | 1.87e-22 |
| UMP1 | YBR173C | ump1Δ | -0.526 | 2.17e-111 |
| GPI10 | YGL142C | gpi10-5001 | -0.523 | 2.90e-40 |
| RPN7 | YPR108W | rpn7-3 | -0.473 | 0.00e+00 |
| RPT2 | YDL007W | rpt2-rf | -0.450 | 2.20e-59 |
| PRE4 | YFR050C | pre4-5001 | -0.449 | 7.87e-46 |
| POC4 | YPL144W | poc4Δ | -0.446 | 1.56e-08 |
| PUP2 | YGR253C | pup2-5001 | -0.417 | 1.98e-37 |
| DPM1 | YPR183W | dpm1-6-supp1 | -0.409 | 1.40e-10 |

**Table S3: Top negative genetic interaction with *DFG10*. Related to Fig 1C.**

| **Gene name** | **Systematic gene name** | **Mutation allele** | **Gene interaction score** | **p-value** |
| --- | --- | --- | --- | --- |
| SSP2 | YOR242C | ssp2Δ | -0.894 | 0.00161 |
| ALG12 | YNR030W | alg12Δ | -0.855 | 0.00003 |
| RET2 | YFR051C | ret2-1 | -0.854 | 0.00038 |
| PMT2 | YAL023C | pmt2Δ | -0.852 | 0.00087 |
| ENV9 | YOR246C | env9Δ | -0.851 | 0.00026 |
| ALG6 | YOR002W | alg6Δ | -0.833 | 0.00031 |
| GPI13 | YLL031C | gpi13-3 | -0.833 | 0.02525 |
| PMT1 | YDL095W | pmt1Δ | -0.803 | 0.00038 |
| ERG9 | YHR190W | erg9-ph | -0.801 | 0.00423 |
| BEM2 | YER155C | bem2Δ | -0.800 | 0.00193 |
| SLT2 | YHR030C | slt2Δ | -0.786 | 0 |
| RGD1 | YBR260C | rgd1Δ | -0.782 | 0.0029 |
| ALG3 | YBL082C | alg3Δ | -0.774 | 0.0001 |
| HAC1 | YFL031W | hac1Δ | -0.755 | 0.00877 |
| CNB1 | YKL190W | cnb1Δ | -0.748 | 0.00408 |
| COP1 | YDL145C | cop1-1 | -0.744 | 0.00456 |
| NUS1 | YDL193W | nus1-ph | -0.737 | 0.00022 |

**Table S4: Top negative genetic interaction with *TDA5 (tda5-*supp-1 (SUM1) allele*)*. Related to Fig 4A.**

| **Systemic gene name** | **Gene name** | **Mutation allele** | **Gene interaction score** | **p-value** |
| --- | --- | --- | --- | --- |
| YMR148W | LDO16 | ldo16Δ | -0.798 | 0.00E+00 |
| YDL193W | NUS1 | nus1-5001 | -0.791 | 4.87E-81 |
| YPL033C | SRL4 | srl4Δ | -0.753 | 1.02E-55 |
| YJL160C | PIR5 | pir5Δ | -0.702 | 4.23E-20 |
| YOR246C | ENV9 | env9Δ | -0.688 | 5.49E-44 |
| YOR242C | SSP2 | ssp2Δ | -0.654 | 1.34E-31 |
| YBR004C | GPI18 | gpi18-5001 | -0.650 | 7.31E-55 |
| YBR070C | ALG14 | alg14-5001 | -0.542 | 0.00E+00 |
| YOR149C | SMP3 | smp3-1 | -0.511 | 9.75E-12 |
| YDR331W | GPI8 | gpi8-5001 | -0.505 | 8.08E-17 |
| YHR188C | GPI16 | gpi16-5001 | -0.503 | 5.16E-12 |
| YMR013C | SEC59 | sec59-ts | -0.501 | 4.36E-82 |
| YDL114W | YDL114W | ydl114wΔ | -0.458 | 8.78E-12 |
| YDR302W | GPI11 | gpi11-5001 | -0.456 | 3.13E-25 |
| YML071C | COG8 | cog8Δ | -0.447 | 2.51E-13 |
| YKR020W | VPS51 | vps51Δ | -0.447 | 1.44E-11 |
| YHL031C | GOS1 | gos1Δ-supp1 | -0.443 | 2.98E-19 |
| YMR272C | SCS7 | scs7Δ | -0.414 | 4.63E-131 |
| YOL135C | MED7 | med7-141 | -0.413 | 2.18E-62 |
| YNL041C | COG6 | cog6Δ | -0.404 | 2.41E-118 |

**Table S5: Top negative genetic interaction with *TDA5 (*tda5*-*supp-2 (HST1) allele*)*.**

| **Gene name** | **Systemic gene name** | **Mutation allele** | **Gene interaction score** | **p-value** |
| --- | --- | --- | --- | --- |
| YOR246C | ENV9 | env9Δ | -0.770 | 6.39E-62 |
| YDL114W | YDL114W | ydl114wΔ | -0.765 | 2.85E-52 |
| YMR148W | LDO16 | ldo16Δ | -0.759 | 0.00E+00 |
| YPL033C | SRL4 | srl4Δ | -0.722 | 8.49E-85 |
| YMR215W | GAS3 | gas3Δ | -0.595 | 0.00E+00 |
| YML066C | SM2 | sma2Δ | -0.539 | 4.80E-130 |
| YPR183W | DPM1 | dpm1-6-supp1 | -0.384 | 0.00E+00 |
| YGL124C | MON1 | mon1Δ | -0.374 | 8.83E-09 |
| YBR002C | RER2 | rer2-5008 | -0.357 | 0.00E+00 |
| YHR107C | CDC12 | cdc12-1 | -0.325 | 2.23E-07 |
| YGL222C | EDC1 | edc1Δ | -0.301 | 8.10E-03 |
| YLR275W | SMD2 | smd2-5005 | -0.252 | 5.65E-22 |
| YJR085C | TMH11 | tmh11Δ | -0.248 | 2.45E-31 |
| YNL055C | POR1 | por1Δ | -0.246 | 1.02E-03 |
| YEL032W | MEM3 | mcm3-1 | -0.238 | 1.24E-02 |
| YCR094W | CDC50 | cdc50Δ | -0.223 | 6.03E-03 |
| YJL089W | SIP4 | sip4Δ | -0.211 | 2.70E-02 |
| YJR076C | CDC11 | cdc11-5 | -0.208 | 2.63E-03 |
| YNR010W | CSE2 | cse2Δ | -0.203 | 2.10E-02 |
| YBL020W | RFT1 | rft1-5028 | -0.200 | 2.67E-03 |

**Table S6. *S. cerevisiae* strains used in this study**

| **Source** | **Strain** | **Relevant genotype/description** |
| --- | --- | --- |
| Brachmann *et al.* 1998 | BY4743 | MATa/MATalfa, his3Δ1/his3Δ1 leu2Δ0/leu2Δ0 met15Δ0/MET15 LYS2/lys2Δ0 ura3Δ0/ura3Δ0 |
| Brachmann *et al.* 1998 | BY4741 | Mat a, his3Δ1 leu2Δ1 met15Δ01 ura3Δ1 |
| Brachmann *et al.* 1998 | BY4742 | Mat alfa ,his3Δ1 leu2Δ1 lys2Δ0 ura3Δ1 |
| This study | DFG10/dfg10 | BY3743 genetic background, Mat a/alfa, HETDip DFG10/dfg10::his3MX^loxP |
| This study | dfg10-1 | BY4741 genetic background, Mat a, dfg10::his3MX6loxP, /pTU-DFG10 |
| This study | dfg10-3 | BY4742 genetic background Mat alfa dfg10::his3MX6loxP, /pTU-DFG10 |
| Dharmacon | clone ID 2502 | BY4741 genetic background, env9::kanMX4 |
| This study | env9 | Derivative of clone ID 2502, BY4741 genetic background, Mat a Env9::natMX4; |
| Dharmacon | clone ID 26029 | BY3743 genetic background, Mat a/alfa HETDip tda5:kanMX4/TDA5/pTU-TDA5 |
| This study | tda5/pTU-TDA5 | Derivative of clone ID 26029, BY4742 genetic background Mat alfa tda5::kanMX4/pTU-TDA5 |
| This study | tda5 | Derivative of clone ID 26029, BY4742 genetic background Mat alfa tda5::kanMX4/ pTU-TDA5-ENV9-YDL114w |
| Dharmacon | clone ID 3811 | BY4741 genetic background, Mat a ydl114w::kanMX4 |
| This study | ydl114w | Derivative of clone ID 3811, BY4741 genetic background, Mat a ydl114w::hphMX3 |
| This study | dfg10-env9 | BY4741 genetic background, dfg10::his3MX6loxP, env9::natMX3/pTU-DFG10 |
| This study | KG801 | Derivative of tda5 and ydl114w Mat alfa tda5:kanMX4, env9::natMX4, his3Δ1 leu2Δ1 met15Δ01 ura3Δ1 lys2Δ0 pTU-TDA5-ENV9-YDL114w |
| This study | KG804 | Derivative of KG801 and ydl114w, Mat alfa tda5:kanMX4, ydl114w::hphMX3, env9::natMX4, env9::natMX6, his3Δ1 leu2Δ1 met15Δ01 ura3Δ1 lys2Δ0 / pTU-TDA5-ENV9-YDL114w |
| Euroscarf | alg3 | BY4741; MATa; his3Δ1; leu2Δ0; met15Δ0; ura3Δ0; Alg3::kanMX4 |

Brachmann CB, Davies A, Cost GJ, Caputo E, Li J, Hieter P, Boeke JD. Designer deletion strains derived from Saccharomyces cerevisiae S288C: a useful set of strains and plasmids for PCR-mediated gene disruption and other applications. *Yeast*. 1998 14:115-32.

**Table S7: Plasmids for *S. cerevisiae* expression system**

| **Source/Addgene ID** | ***S.cerevisiae* plasmids** | **Description** |
| --- | --- | --- |
| Park *et al.* 2014/#203163 | pKG-GW1 | *LEU2* marker, 2μ origin of replication, Gateway Cloning ready, TDH3 yeast promoter |
| Galosi *et al.* 2022/#203176 | pTL | Centromeric plasmid, *LEU2* marker, Gateway Cloning ready |
| Van Mullem *et al.* 2003 | pVV208 | Centromeric plasmid, *URA3* marker, Gateway Cloning ready |
| This study/#203179 | pTU | Derivative of pVV208 lacking tetracycline response element, URA3 marker, Gateway Cloning ready |
| This study/#203178 | pTM | Centromeric plasmid, *MET17* marker, Gateway Cloning ready |
| This study/#203183 | pKG-GW1-DHRSX | Derivative of pKG-GW1 expressing DHRSX |
| This study/#203184 | pKG-GW1-TSC13 | Derivative of pKG-GW1 expressing TSC13 |
| This study/#203185 | pKG-GW1-SPS19 | Derivative of pKG-GW1 expressing SPS13 |
| This study/#203186 | pKG-GW1-DFG10 | Derivative of pKG-GW1 expressing DFG10 |
| This study/#203187 | pKG-GW1-TDA5 | Derivative of pKG-GW1 expressing TDA5 |
| This study/#203188 | pKG-GW1-YDL114w | Derivative of pKG-GW1 expressing YDL114w |
| This study/#203189 | pKG-GW1-ENV9 | Derivative of pKG-GW1 expressing ENV9 |
| This study/#203190 | pTU-TDA5 | Derivative of pTU expressing TDA5 under native promoter |
| This study/#203191 | pTU-TDA5-ENV9-YDL114w | Derivative of pTU expressing TDA5, ENV9 and YDL114w under native promoters |
| This study/#203202 | pTU-DFG10 | Derivative of pTU expressing DFG10 under native promoter |
| This study/#203203 | pTL-DFG10 | Derivative of pTU expressing DFG10 under native promoter |
| This study/#203192 | pTL-TDA5 | Derivative of pTL expressing TDA5 under native promoter |
| This study/#203194 | pTL-ENV9 | Derivative of pTL expressing ENV9 under native promoter |
| This study/#203196 | pTL-YDL114w | Derivative of pTL expressing YDL114w under native promoter |
| This study/#203204 | pTL-TDA5/ENV9 | Derivative of pTL expressing TDA5 and ENV9 under native promoters |
| This study/#203205 | pTL- TDA5/YDL114w | Derivative of pTL expressing TDA5 and YDL114w under native promoters |
| This study/#203206 | pTL-YDL114w/ENV9 | Derivative of pTL expressing YDL114w and EV9 under native promoters |
| This study/#203208 | pTL- TDA5/YDL114w/ENV9 | Derivative of pTL expressing TDA5, YDL114w and ENV9 under native promoters |
| This study/#203199 | pTM-DFG10-6FLAG | Derivative of pTM expressing C-terminally FLAG tagged DFG10 under native promoter |

Galosi S, Edani BH, Martinelli S, Hansikova H, Eklund EA, Caputi C, Masuelli L, Corsten-Janssen N, Srour M, Oegema R, Bosch DGM, Ellis CA, Amlie-Wolf L, Accogli A, Atallah I, Averdunk L, Barañano KW, Bei R, Bagnasco I, Brusco A, Demarest S, Alaix AS, Di Bonaventura C, Distelmaier F, Elmslie F, Gan-Or Z, Good JM, Gripp K, Kamsteeg EJ, Macnamara E, Marcelis C, Mercier N, Peeden J, Pizzi S, Pannone L, Shinawi M, Toro C, Verbeek NE, Venkateswaran S, Wheeler PG, Zdrazilova L, Zhang R, Zorzi G, Guerrini R, Sessa WC, Lefeber DJ, Tartaglia M, Hamdan FF, Grabińska KA, Leuzzi V. De novo DHDDS variants cause a neurodevelopmental and neurodegenerative disorder with myoclonus. Brain. 2022 145:208-223

Park EJ, Grabińska KA, Guan Z, Stránecký V, Hartmannová H, Hodaňová K, Barešová V, Sovová J, Jozsef L, Ondrušková N, Hansíková H, Honzík T, Zeman J, Hůlková H, Wen R, Kmoch S, Sessa WC. Mutation of Nogo-B receptor, a subunit of cis-prenyltransferase, causes a congenital disorder of glycosylation. Cell Metab. 2014 20:448-57

Van Mullem V, Wery M, De Bolle X, Vandenhaute J. Construction of a set of Saccharomyces cerevisiae vectors designed for recombinational cloning. Yeast. 2003 20:739-46.

HiFi DNA Assembly method (NEBuilder®, NEB) was used to construct expression vectors. Yeast genomic DNA was used as template to amplify yeast genes.

**Insert sequences:**

P_TDA5_-TDA5 (*TDA5* promoter lower case, ORF upper case)

aatctagtttttttgccttggtttatgatgaagccgtagtgaaaataaactgctacggtgagatgtctgattggagaaaggatattctcttacttgatttttgttgtaccggcgcgagcttccatggtaatcatttgattctcgttggcgacaatttgatacagatttacgacttaaaaaatgttgagcagaacctaggcgaattggttcccgtccagatcataaaaggaaagaagataaagctggctagttcagaaaggagagagaaaacaattcttgtgttaagtcatccaaatatacttaatagacaattattagtcgcgtgcaatcctgtagcaatggcagatcatcaataatataagatatagacacttttaatggtaaatatcgttgtaataatataagccaacaagaatatagtccagtaattatgctatttgcagtaactcacatcattattctctctgttcatataatcctgaattacaggttaccctcggaagtgcatttagccgtataacaagtagctctttaaagccagtagaacagatacattaatatgtgataatacggctgtgaggaaaaggtgaagttctggtgATGAATATAGACTGTTTGTGTCGTTGGGTAGTTCTGCCACTGCTGCGCTATCCGCTTCTGGTAGCTCTTGTATTGAGATGGTCTCTAAGCGACTCGATAAGCATTTGTCTAACAATCTATACACTACTGATAAATGCATTCCTCATTGCTAATTCCTATACAAAGCGGTCGGGACAGGTCGCCTGGAAGAGCCTTCGTGAATTTAAGAATGGCATAGTTCTTATTACTGGAGGAAGTAAAGGGCTTGGACGGGCCATAGTATCTCAACTCCTACAAGACTACAGTAACCTGACCATCTTGAATGTCGACATATGCCCATCATCAGTGAGGAATACTCGCGTGAAGGATCTCATCTGCGACCTGAGTGATGATGAAGAAGTTGCAGCACTACTGAATTTGTTGAAAAGAAAGTATAAAAATGAAATCCGATTGATTGTGAACAATGCTGGAGTAAGGGCCAACTTCACCGGATTCAACGGCATGGAACGGGATAATCTTGATAAGATTTTTAAGATAAACACATTTGCACCACTTCAGTTCATTCAAGAACTGGCCCCTAGTAGACACTCAACTAGACAGTGTTATATTGTCAATATCGCAAGCATTTTGGGCATATTGACGCCGGCCAAAGTAGCAGCTTATGCGGCGAGCAAAGCAGCATTGATAGCCTTTCATCAGTCGTACAGTTTTGAATTGCAAAACGAAGGCGTAAGAAATATCAGAACACTATTAGTGACCCCAGGGCAACTGAATACGGAAATGTTTGCAGGATTCAAGCCTCCTCGCCAGTTCTTTGCCCCTGTAATAGACATTACTACTCTAGCTGCCAAGATAGTCCGTTATTGTGAGCTGGGTCAGAGGGGACAGCTAAATGAACCCTTTTATTGTAGTTTTGCTCATCTTTTGATGTGCGTACCTTATTCGCTACAACGCATTGTAAGAAGCTTCTCTCGCATAGATTGCTGCCTCCCGGACGAGTAG

P_ENV9_-ENV9 (*ENV9* promoter lower case, ORF upper case)

ggtagaggtaagaaagaataaaccaaaatgtaagctaggtgagggaaatattttggagcttaaatatctcgaacgacactcactctcgtcagaaaaatctgtcgagtaactcagaccacctccgatagcataccttgtttagtttcataatcggcagagacgtcattcctgcacagaacgccaaaattcgtggcccccgcggtacctgaatcagggccaagaattttggcgtttcgtgttgttgacatattccgaaaaaggtattgagccacaggtctttcgcgtcttctaagccccaagataaagaaaatgccgccctgcttaatcactgtttttttttggaaaaccctgaagtatagaagtagggcattgaaagaaaagatatgcgggtaagatatgaaaaaattgaggaaatgagacccaagtgtacatgggtatatataaacaaatcattatggtgcatgatgcatcacttacaatgcacacacactattgtctttgtagaaacggggaagacaatggttgtaggcctgattggtattagaggcatccgcagtacgagacatacgccaaaggcagaATGTTAGACCCACGAATATTGCCATACTACGACCCGGCTGTGGAGAGGAAGATTGCTGTAGTAACAGGCGGAAATACGGGTATTGGGTGGTATACTGTCTTGCATTTGTATTTGCATGGGTTTGTCGTTTATATTTGTGGGAGAAACTCTCACAAGATTTCGAAAGCAATCCAGGAGATACTGGCAGAAGCAAAGAAGAGGTGCCATGAAGATGACGACGGTTCAAGCCCAGGCGCGGGCCCAGGTCCAAGCATTCAGCGTCTAGGGTCACTGCACTATATCCATTTGGATTTGACAGACTTAAAATGTGTGGAGAGAGCGGCACTTAAAATTCTCAAGCTGGAAGACCACATAGACGTGCTTGTTAACAATGCAGGGATTATGGCGGTGCCCTTAGAAATGACGAAGGACGGATTTGAAGTGCAATTGCAGACTAACTACATTTCGCACTTCATCTTCACGATGAGATTATTGCCTTTACTGCGCCATTGTCGTGGCAGGATCATTTCCCTGTCCTCGATAGGCCATCATCTAGAGTTCATGTACTGGAAACTGAGCAAGACGTGGGATTACAAACCTAATATGCTTTTCACATGGTTTAGGTACGCGATGAGTAAAACCGCGCTAATCCAATGCACGAAGATGTTGGCCATCAAATACCCTGACGTTCTTTGTCTCTCCGTTCATCCGGGTCTGGTGATGAACACAAACTTATTCAGTTATTGGACAAGGTTACCCATCGTCGGTATTTTCTTTTGGCTGTTGTTCCAGGTCGTAGGGTTCTTTTTCGGCGTATCGAACGAACAAGGTTCACTAGCTTCTTTGAAGTGTGCATTGGACCCGAATTTATCTGTCGAAAAAGATAACGGGAAGTACTTCACCACGGGGGGTAAAGAATCTAAATCGAGCTACGTTTCTAACAATGTCGACGAAGCGGCATCGACTTGGATCTGGACCGTTCATCAACTAAGAGACCGTGGTTTCGATATATAA

P_YDL114w_-YDL114w (*YDL114w* promoter lower case, ORF upper case)

caaacgcatttttgttatatcggtagaataacaacacatacataatgcctgcacagaatcctcatcccttcttctctttactctgataaactcaggtgccgttgtggtactaatagtgctcattcccgaaatcttgtatgcctacctcaacaaaagctatacctcagagagcagttttatataccatctcttgttctcatcaataacttgtcttgagcattacacaatttaaatcccctgtgccgacacaaacaagagataacccggaatcaatattatctgattgggctcctaaatatcgcatctgtggacttgaatccacatgctgcgaacttatttgaagcaacaattacttctgttggaagcgtggtagtgccaaatcagacttgctctttagttttcttatgttaacgctttactgcctttttcttctatatttttgaacgtcaatatgtagcgggcgtcagcagaggaacagaatttgtgacacttcttgaatgaacatttctcatatctgaaaccgcatcatcaaaaggacacctcaatagagattcagtgacatatttaacatagttttatgggtaagaaatattattaattATGGTACGCAAGAATAAAATAAATAGAGCGAGTGGAACTACAAAACACCTCAAGGATTTTCCATCAGTCATCCTAAGTCTGCCCAGCTACAACCCTTCCATCCTTTCTAAAAATGCAACTGCACTAATAACCGGTGGTTCTAGTGGACTCGGATTTGAACTTGCAAAGGAACTTTCCAGAAGAATTAATAAAGTTATTGTGGCAGATATTCAATCATTTCCTACTTTTGCTCAGGTAGAATATAATAACATTTTCTACTATCAATGCGACATAACAAGTCTAGATGAAATTAAAAATTTGAAAAAAGCTATTGAAAGAGATCATGGCAACATTAATATAATTATAAATAATGCTGGTGTTGCTCACATCAAAAAGTTAGAACATATGACAAACAAGGAGGTAGAACAATTGATCGACATAAACTTAATAGGTGCTTACCGAATCATCAGTACATTTGCAGAAGATATGATAGACAATAGGGAAGGCTTTATAATTAACATTGCTTCAGTCCTCGGAGAACTGACTCCTGCAAGGCTGACATCGTATGGTGCATCAAAAGGAGCAATGATTGGTTTTCACAAGTGTATGAGCAGGCATTTCAGAAGTTTGTCCACGGAATGTAATAAAACTGGAATAAAAACACTACTGGTGTGCCCGGGAAAAATCAAAACTAATATGTTTATTGATGTCCCTACACCATCTAAGTTATTGGCTCCTGACATCATACCTTCCCAACTGGCACTAGCGATAATCTCTGCGATGGAGCATAATCATCTGCAAACACTAAATGCCCCATATTATGTGAACCTAGTTCCCTTTTTCAAAACTTTGAGTTGGCCATATAGGCATCTCTTAAAACATTTCAGTGGAATGGACCATGTAACGTCAACCAGCCCGCGAGCAATAAATCCGAAGAGAAGTGCATAA

P_DFG10_-DFG10 (*DFG10* promoter lower case, ORF upper case)

ttgctcataacaggtgtcacaatttgcacgaagttcttgagtgattttttttacagtaactctcgatacgccaaggttggaggtatttctttgcaagagttaaaccatttagaattacaattcctgatactatgtgatttcaagttattggtttcagtcgaagaaatgcagaaatatgcaaacttgttatataaattctggaacgaccagtagaatgctgtcctgcatattattcatgcccttcccttaccgttactcgtattatacatatcaatttctgatttacgtaacgcgtacatatcggttatatttaaataatttaatattccttttaaatcagtggtttttcatattagtagcatttctcctgaaatgatagctattgcaatagaatgaaaaaaaaaactttttgcccaaataaatcataatctaccggtaaagagaaattcggtccttgttctattcagtattatgctttctcttttttttccaaaatgagaattaaggacgtttcgaaaaagtaaaaatataccaaatgtggaatataaataggtttatcggcggaccgatctggcttaaaacgtagaatctaccagtccaATGTACTTTGATGAAGAACAATTGCTAAAATATACTATATATGCCTATAGATTATCCTTTTTTGTAGGCATTTGCTCACTTTTCATAGCAAAAAGTTGTCTACCAGAATTTCTTCAATATGGTAAAACCTACCGGCCCAAAGAGAATTCAAAGTACTCAAGCATTTTAGAACGAATCAAGAAGTTCACAGTTCCAAAGGCGTATTTTTCCCATTTTTACTATTTGGCTACCTTTCTATCCTTAGTCACCTTATATTTCTATCCTAAATTCCCCATCGTTTGGATCATATTTGGACACTCATTGCGCCGACTTTATGAAACGCTTTATGTACTACATTATACAAGCAATTCTAGGATGAATTGGTCCCATTATCTAGTCGGTATATGGTTCTATTCCGTACTCTTGTTAATTCTTAATATATCACTGTACAAGAACTCCATTCCAAATACGTTAAACATGAATGCTTTCATCATATTCTGCATAGCATCTTGGGATCAGTACAAAAATCATGTTATTCTGGCCAATCTGGTTAAATATTCGCTGCCAACAGGAAGGCTTTTCAGGTTGGTATGCTGTCCTCATTATCTCGATGAAATAATCATTTATTCTACTCTGTTGCCCTATGAACAAGAATTTTACCTAACACTAGTTTGGGTAATCACAAGTTTGACTATATCCGCATTGGAAACAAAAAATTATTACAGGCACAAATTTAAAGACAATCACGTAGCCCCCTACGCCATAATACCTTTTATAATCTAG

P_DFG10_-DFG10-6FLAG (*DFG10* promoter lower case, ORF upper case, 6FLAG italic, lower case)

ttgctcataacaggtgtcacaatttgcacgaagttcttgagtgattttttttacagtaactctcgatacgccaaggttggaggtatttctttgcaagagttaaaccatttagaattacaattcctgatactatgtgatttcaagttattggtttcagtcgaagaaatgcagaaatatgcaaacttgttatataaattctggaacgaccagtagaatgctgtcctgcatattattcatgcccttcccttaccgttactcgtattatacatatcaatttctgatttacgtaacgcgtacatatcggttatatttaaataatttaatattccttttaaatcagtggtttttcatattagtagcatttctcctgaaatgatagctattgcaatagaatgaaaaaaaaaactttttgcccaaataaatcataatctaccggtaaagagaaattcggtccttgttctattcagtattatgctttctcttttttttccaaaatgagaattaaggacgtttcgaaaaagtaaaaatataccaaatgtggaatataaataggtttatcggcggaccgatctggcttaaaacgtagaatctaccagtccaATGTACTTTGATGAAGAACAATTGCTAAAATATACTATATATGCCTATAGATTATCCTTTTTTGTAGGCATTTGCTCACTTTTCATAGCAAAAAGTTGTCTACCAGAATTTCTTCAATATGGTAAAACCTACCGGCCCAAAGAGAATTCAAAGTACTCAAGCATTTTAGAACGAATCAAGAAGTTCACAGTTCCAAAGGCGTATTTTTCCCATTTTTACTATTTGGCTACCTTTCTATCCTTAGTCACCTTATATTTCTATCCTAAATTCCCCATCGTTTGGATCATATTTGGACACTCATTGCGCCGACTTTATGAAACGCTTTATGTACTACATTATACAAGCAATTCTAGGATGAATTGGTCCCATTATCTAGTCGGTATATGGTTCTATTCCGTACTCTTGTTAATTCTTAATATATCACTGTACAAGAACTCCATTCCAAATACGTTAAACATGAATGCTTTCATCATATTCTGCATAGCATCTTGGGATCAGTACAAAAATCATGTTATTCTGGCCAATCTGGTTAAATATTCGCTGCCAACAGGAAGGCTTTTCAGGTTGGTATGCTGTCCTCATTATCTCGATGAAATAATCATTTATTCTACTCTGTTGCCCTATGAACAAGAATTTTACCTAACACTAGTTTGGGTAATCACAAGTTTGACTATATCCGCATTGGAAACAAAAAATTATTACAGGCACAAATTTAAAGACAATCACGTAGCCCCCTACGCCATAATACCTTTTATAATCgga*gactacaaagaccatgacggtgattataaagatcatgacatcgattacaaggatgacgatgacaagggagactataaggaccacgacggggactacaaggaccacgatatcgactataaagacgatgacgataaa*tag

DHRSX (restriction sites and attB1/attB2 lower case; ORF upper case)

acaagtttgtacaaaaaagcaggctccgaattcgcccttcatATGTCGCCATTGTCTGCGGCGCGGGCGGCCCTGCGGGTCTACGCGGTAGGCGCCGCGGTGATCCTGGCGCAGCTGCTGCGGCGCTGCCGCGGGGGCTTCCTGGAGCCAGTTTTCCCCCCACGACCTGACCGTGTCGCTATAGTGACGGGAGGGACAGATGGCATTGGCTATTCTACAGCGAAGCATCTGGCGAGACTTGGCATGCATGTTATCATAGCTGGAAATAATGACAGCAAAGCCAAACAAGTTGTAAGCAAAATAAAAGAAGAAACCTTGAACGACAAAGTGGAATTTTTATACTGTGACTTGGCTTCCATGACTTCCATCCGGCAGTTTGTGCAGAAGTTCAAGATGAAGAAGATTCCTCTCCATGTCCTGATCAACAATGCTGGGGTGATGATGGTCCCTCAGAGGAAAACCAGAGATGGATTCGAAGAACATTTCGGCCTGAACTACCTAGGGCACTTCCTGCTGACCAACCTTCTCTTGGATACGCTGAAAGAGTCTGGGTCCCCTGGCCACAGTGCGAGGGTGGTCACCGTCTCCTCTGCCACCCATTACGTCGCTGAGCTGAACATGGATGACCTTCAGAGCAGTGCCTGCTACTCACCCCACGCAGCCTACGCCCAGAGCAAGCTGGCCCTTGTCCTGTTCACCTACCACCTCCAGCGGCTGCTGGCGGCTGAGGGAAGCCACGTGACCGCCAACGTGGTGGACCCCGGGGTGGTCAACACGGACCTCTACAAGCACGTGTTCTGGGCCACCCGTCTGGCGAAGAAGCTTCTCGGCTGGTTGCTTTTCAAGACCCCCGATGAAGGAGCGTGGACTTCCATCTACGCAGCAGTCACCCCAGAGCTGGAAGGAGTTGGTGGCCGTTACCTATACAACGAGAAAGAGACCAAGTCCCTCCACGTCACCTACAACCAGAAACTGCAGCAGCAGCTGTGGTCTAAGAGTTGTGAGATGACTGGGGTCCTTGATGTGACCCTGTGAgcggccgcaagggcgaattcgacccagctttcttgtacaaagtggt

TSC13 (restriction sites and attB1/attB2 lower case; ORF upper case)

acaagtttgtacaaaaaagcaggctccgaattcgcccttctcgagATGCCTATCACCATAAAAAGCCGCTCTAAAGGGTTAAGGGACACTGAAATTGACTTATCCAAAAAGCCTACTTTAGATGATGTTTTGAAAAAAATCTCTGCTAATAACCACAATATCAGCAAGTACAGGATAAGATTAACCTACAAAAAGGAATCTAAACAAGTTCCGGTTATTTCAGAATCGTTTTTTCAAGAAGAGGCTGATGACTCAATGGAATTCTTCATCAAAGATTTGGGTCCCCAAATTTCATGGAGATTAGTCTTCTTTTGTGAGTATTTGGGTCCAGTCTTGGTTCACTCCCTTTTTTATTATCTATCTACCATTCCCACAGTTGTTGATAGATGGCACAGTGCTAGCTCCGACTATAATCCATTTTTAAACAGGGTTGCATATTTTTTAATTTTAGGACATTATGGAAAGAGATTATTTGAAACCTTATTTGTTCACCAATTCTCTTTAGCTACTATGCCAATTTTCAACCTGTTCAAAAATTGTTTCCATTACTGGGTTCTAAGCGGTCTCATTTCATTCGGTTACTTTGGCTACGGCTTCCCCTTTGGGAATGCTAAGTTATTCAAATACTATTCATATTTGAAATTGGATGACTTGAGTACATTAATTGGTCTTTTCGTGCTTTCAGAACTATGGAACTTTTATTGCCACATTAAATTGCGCCTATGGGGTGACTATCAAAAGAAGCATGGTAACGCTAAGATCCGTGTCCCATTGAATCAAGGTATTTTCAATCTTTTTGTTGCTCCCAACTATACTTTTGAAGTTTGGTCTTGGATTTGGTTTACTTTTGTGTTCAAGTTCAATTTATTTGCCGTTTTATTTTTGACTGTTTCAACAGCTCAAATGTACGCATGGGCTCAAAAGAAAAACAAAAAGTATCATACCAGAAGAGCATTCTTGATTCCATTTGTATTTTGAgcggccgcaagggcgaattcgacccagctttcttgtacaaagtggt

SPS19 (restriction sites and attB1/attB2 lower case; ORF upper case)

acaagtttgtacaaaaaagcaggctccgaattcgcccttctcgagATGAATACAGCAAACACTTTGGACGGCAAATTCGTTACTGAGGGTTCTTGGAGACCTGATTTATTTAAGGGTAAAGTGGCATTTGTCACTGGTGGAGCTGGCACGATATGTCGGGTACAGACAGAAGCCTTGGTTCTTCTTGGCTGTAAGGCAGCTATTGTCGGCAGGGACCAAGAAAGAACAGAACAGGCAGCAAAAGGTATTTCGCAGTTAGCAAAGGACAAAGATGCAGTCTTGGCTATTGCAAACGTCGATGTTCGCAATTTTGAACAAGTGGAGAATGCGGTTAAAAAGACCGTAGAGAAGTTCGGTAAGATCGATTTCGTCATTGCCGGTGCTGCTGGAAATTTTGTGTGCGATTTTGCGAACCTCTCTCCAAACGCCTTCAAATCTGTTGTTGACATAGATTTACTAGGAAGTTTTAATACCGCCAAGGCATGCTTGAAGGAGTTAAAAAAGTCAAAAGGTTCCATCCTTTTCGTCTCTGCCACTTTCCATTATTATGGTGTGCCGTTTCAAGGACATGTGGGCGCTGCTAAAGCGGGAATAGACGCATTGGCAAAGAATTTAGCGGTTGAGTTGGGGCCTCTGGGTATACGTTCGAATTGCATTGCTCCAGGTGCTATCGATAATACGGAAGGCTTGAAGAGATTAGCCGGCAAAAAATATAAGGAAAAGGCCTTGGCAAAGATACCATTACAAAGGCTTGGATCTACACGGGATATAGCCGAATCTACTGTTTATATTTTTTCACCCGCCGCATCGTATGTCACCGGTACCGTCTTAGTTGTTGACGGTGGTATGTGGCATTTAGGGACCTATTTTGGACATGAGTTGTATCCAGAAGCCTTAATAAAGAGTATGACATCTAAATTATAAgcggccgcaagggcgaattcgacccagctttcttgtacaaagtggt

DFG10 (restriction sites and attB1/attB2 lower case; ORF upper case)

acaagtttgtacaaaaaagcaggctccgaattcgcccttctcgagATGTACTTTGATGAAGAACAATTGCTAAAATATACTATATATGCCTATAGATTATCCTTTTTTGTAGGCATTTGCTCACTTTTCATAGCAAAAAGTTGTCTACCAGAATTTCTTCAATATGGTAAAACCTACCGGCCCAAAGAGAATTCAAAGTACTCAAGCATTTTAGAACGAATCAAGAAGTTCACAGTTCCAAAGGCGTATTTTTCCCATTTTTACTATTTGGCTACCTTTCTATCCTTAGTCACCTTATATTTCTATCCTAAATTCCCCATCGTTTGGATCATATTTGGACACTCATTGCGCCGACTTTATGAAACGCTTTATGTACTACATTATACAAGCAATTCTAGGATGAATTGGTCCCATTATCTAGTCGGTATATGGTTCTATTCCGTACTCTTGTTAATTCTTAATATATCACTGTACAAGAACTCCATTCCAAATACGTTAAACATGAATGCTTTCATCATATTCTGCATAGCATCTTGGGATCAGTACAAAAATCATGTTATTCTGGCCAATCTGGTTAAATATTCGCTGCCAACAGGAAGGCTTTTCAGGTTGGTATGCTGTCCTCATTATCTCGATGAAATAATCATTTATTCTACTCTGTTGCCCTATGAACAAGAATTTTACCTAACACTAGTTTGGGTAATCACAAGTTTGACTATATCCGCATTGGAAACAAAAAATTATTACAGGCACAAATTTAAAGACAATCACGTAGCCCCCTACGCCATAATACCTTTTATAATCTAGgcggccgcaagggcgaattcgacccagctttcttgtacaaagtggt

YDL114w (restriction sites and attB1/attB2 region italic; ORF upper case)

acaagtttgtacaaaaaagcaggctccgaattcgcccttctcgagATGGTACGCAAGAATAAAATAAATAGAGCGAGTGGAACTACAAAACACCTCAAGGATTTTCCATCAGTCATCCTAAGTCTGCCCAGCTACAACCCTTCCATCCTTTCTAAAAATGCAACTGCACTAATAACCGGTGGTTCTAGTGGACTCGGATTTGAACTTGCAAAGGAACTTTCCAGAAGAATTAATAAAGTTATTGTGGCAGATATTCAATCATTTCCTACTTTTGCTCAGGTAGAATATAATAACATTTTCTACTATCAATGCGACATAACAAGTCTAGATGAAATTAAAAATTTGAAAAAAGCTATTGAAAGAGATCATGGCAACATTAATATAATTATAAATAATGCTGGTGTTGCTCACATCAAAAAGTTAGAACATATGACAAACAAGGAGGTAGAACAATTGATCGACATAAACTTAATAGGTGCTTACCGAATCATCAGTACATTTGCAGAAGATATGATAGACAATAGGGAAGGCTTTATAATTAACATTGCTTCAGTCCTCGGAGAACTGACTCCTGCAAGGCTGACATCGTATGGTGCATCAAAAGGAGCAATGATTGGTTTTCACAAGTGTATGAGCAGGCATTTCAGAAGTTTGTCCACGGAATGTAATAAAACTGGAATAAAAACACTACTGGTGTGCCCGGGAAAAATCAAAACTAATATGTTTATTGATGTCCCTACACCATCTAAGTTATTGGCTCCTGACATCATACCTTCCCAACTGGCACTAGCGATAATCTCTGCGATGGAGCATAATCATCTGCAAACACTAAATGCCCCATATTATGTGAACCTAGTTCCCTTTTTCAAAACTTTGAGTTGGCCATATAGGCATCTCTTAAAACATTTCAGTGGAATGGACCATGTAACGTCAACCAGCCCGCGAGCAATAAATCCGAAGAGAAGTGCATAAgcggccgcaagggcgaattcgacccagctttcttgtacaaagtggt

ENV9 (restriction sites and attB1/attB2 region italic; ORF upper case)

acaagtttgtacaaaaaagcaggctccgaattcgcccttcatATGTTAGACCCACGAATATTGCCATACTACGACCCGGCTGTGGAGAGGAAGATTGCTGTAGTAACAGGCGGAAATACGGGTATTGGGTGGTATACTGTCTTGCATTTGTATTTGCATGGGTTTGTCGTTTATATTTGTGGGAGAAACTCTCACAAGATTTCGAAAGCAATCCAGGAGATACTGGCAGAAGCAAAGAAGAGGTGCCATGAAGATGACGACGGTTCAAGCCCAGGCGCGGGCCCAGGTCCAAGCATTCAGCGTCTAGGGTCACTGCACTATATCCATTTGGATTTGACAGACTTAAAATGTGTGGAGAGAGCGGCACTTAAAATTCTCAAGCTGGAAGACCACATAGACGTGCTTGTTAACAATGCAGGGATTATGGCGGTGCCCTTAGAAATGACGAAGGACGGATTTGAAGTGCAATTGCAGACTAACTACATTTCGCACTTCATCTTCACGATGAGATTATTGCCTTTACTGCGCCATTGTCGTGGCAGGATCATTTCCCTGTCCTCGATAGGCCATCATCTAGAGTTCATGTACTGGAAACTGAGCAAGACGTGGGATTACAAACCTAATATGCTTTTCACATGGTTTAGGTACGCGATGAGTAAAACCGCGCTAATCCAATGCACGAAGATGTTGGCCATCAAATACCCTGACGTTCTTTGTCTCTCCGTTCATCCGGGTCTGGTGATGAACACAAACTTATTCAGTTATTGGACAAGGTTACCCATCGTCGGTATTTTCTTTTGGCTGTTGTTCCAGGTCGTAGGGTTCTTTTTCGGCGTATCGAACGAACAAGGTTCACTAGCTTCTTTGAAGTGTGCATTGGACCCGAATTTATCTGTCGAAAAAGATAACGGGAAGTACTTCACCACGGGGGGTAAAGAATCTAAATCGAGCTACGTTTCTAACAATGTCGACGAAGCGGCATCGACTTGGATCTGGACCGTTCATCAACTAAGAGACCGTGGTTTCGATATATAAgcggccgcaagggcgaattcgacccagctttcttgtacaaagtggt

TDA5 (restriction sites and attB1/attB2 region italic; ORF upper case)

acaagtttgtacaaaaaagcaggctccgaattcgcccttctcgagATGAATATAGACTGTTTGTGTCGTTGGGTAGTTCTGCCACTGCTGCGCTATCCGCTTCTGGTAGCTCTTGTATTGAGATGGTCTCTAAGCGACTCGATAAGCATTTGTCTAACAATCTATACACTACTGATAAATGCATTCCTCATTGCTAATTCCTATACAAAGCGGTCGGGACAGGTCGCCTGGAAGAGCCTTCGTGAATTTAAGAATGGCATAGTTCTTATTACTGGAGGAAGTAAAGGGCTTGGACGGGCCATAGTATCTCAACTCCTACAAGACTACAGTAACCTGACCATCTTGAATGTCGACATATGCCCATCATCAGTGAGGAATACTCGCGTGAAGGATCTCATCTGCGACCTGAGTGATGATGAAGAAGTTGCAGCACTACTGAATTTGTTGAAAAGAAAGTATAAAAATGAAATCCGATTGATTGTGAACAATGCTGGAGTAAGGGCCAACTTCACCGGATTCAACGGCATGGAACGGGATAATCTTGATAAGATTTTTAAGATAAACACATTTGCACCACTTCAGTTCATTCAAGAACTGGCCCCTAGTAGACACTCAACTAGACAGTGTTATATTGTCAATATCGCAAGCATTTTGGGCATATTGACGCCGGCCAAAGTAGCAGCTTATGCGGCGAGCAAAGCAGCATTGATAGCCTTTCATCAGTCGTACAGTTTTGAATTGCAAAACGAAGGCGTAAGAAATATCAGAACACTATTAGTGACCCCAGGGCAACTGAATACGGAAATGTTTGCAGGATTCAAGCCTCCTCGCCAGTTCTTTGCCCCTGTAATAGACATTACTACTCTAGCTGCCAAGATAGTCCGTTATTGTGAGCTGGGTCAGAGGGGACAGCTAAATGAACCCTTTTATTGTAGTTTTGCTCATCTTTTGATGTGCGTACCTTATTCGCTACAACGCATTGTAAGAAGCTTCTCTCGCATAGATTGCTGCCTCCCGGACGAGTAGgcggccgcaagggcgaattcgacccagctttcttgtacaaagtggt
